# A Synthetic Protein Secretion System for Living Bacterial Therapeutics

**DOI:** 10.1101/2023.06.14.544856

**Authors:** Recep Erdem Ahan, Cemile Elif Ozcelik, Irem Niran Cagil, Urartu Ozgur Safak Seker

## Abstract

Bacteria species can thrive and colonize different parts of the human body. Those naturally residing at disease sites e.g., tumors and gut can be designed for targeted protein delivery which can provide better clinical profiles for protein-based therapies. Therefore, a generalizable, efficient, and safe protein secretion system would a be valuable tool to engineer therapeutically active microbes, especially for gram-negative species due to the presence of the second cell wall. Here, we propose an approach called iLOM-SS, an acronym for **i**nducible **<u>L</u>**eaky **<u>O</u>**uter **<u>M</u>**embrane based **<u>S</u>**ecretion **<u>S</u>**ystem, to secrete proteins in gram-negative bacteria (GNB). In iLOM-SS, the outer membrane of GNB is made permeable by transient suppression of structural protein(s) to enable free diffusion of cargo proteins expressed at the periplasm. To validate this approach, an iLOM-SS is constructed in *Escherichia coli* Nissle 1917 (EcN) strain. Proteins including enzymes and a human cytokine were proven to be secreted with iLOM-SS by EcN *in vitro*. Further characterizations of iLOM-SS in ECN showed that fast and titratable secretion, a stop switch design for secretion, and functional implementation of the secretion system in different genetic circuit architectures were possible. We foresee that this work will pave the way for designing GNB to secrete proteins for diverse arrays of applications including but not limited to the development of sentinel cells for therapeutic purposes.

## MAIN

Human body is colonized at the very beginning of life by different kinds of microbes whose function and abundance are strongly linked to various pathological conditions.^1^ Due to their profound effect on human health and constant interaction with the body, human-associated microbes are engineered with synthetic biology methodologies to sense diseases related pathological conditions and respond to detected signals for therapeutical purposes.^2–7^ Those engineered microbes offer a platform to address unmet medical needs for many conditions because embedded complex actuator circuits in microbes, such as genetic logic gates, enable targeted and spatiotemporal delivery of multiple payloads.^8–10^

Protein-based payloads, especially protein based biotech drugs, necessitate protein secretion through cell membrane, which is a troublesome process to implement in GNB because of the outer cell wall. For extracellular protein translocation, GNB possess different protein secretion systems which can carry proteins across the membranes by either one- or two-step process.

Briefly, intracellular expressed proteins are directly secreted to extracellular matrix in Type I, III, IV, and VI secretion systems whereas proteins are intermediately translocated to the periplasm by Sec or Tat translocon in Type II, and V secretion systems as well as in curli biogenesis system.^11,12^

Repurposing native secretion systems for extracellular payload translocation is associated with many drawbacks that hamper the development of protein secreting GNB based living bacterial therapeutics. Firstly, recombinant expression of the secretory system itself is required for all systems, many of which have attribution to pathogenicity.^13–16^ In addition, construction, and utilization of these nanomachines put a significant amount of burden on cellular metabolism.^17–19^ Secondly, secretion systems have narrow substrate compatibility; in other words, a small subset of payloads can be secreted by using these systems.^20–22^ Even if a payload protein is secreted to extracellular space, the secreted protein will carry an additional secretion tag that can affect the – protein’s function or potentially cause unwanted inflammatory responses.^17,18,23^ Lastly, deployment of the optimized secretion system from one species to another generally is not possible as systems require many additional elements such, as chaperone proteins, for proper function.^20,22^ Thus achieving production of secretory proteins in non-model Gram-negative bacteria with characterized secretion systems often requires cumbersome optimizations that heavily rely on trial-and-error methods.

To overcome these hurdles related to native secretion systems, many researchers developed unusual ways to release proteins to extracellular matrix in GNB. Din et al. demonstrated that intracellularly expressed recombinant proteins can be released to the extracellular environment with an expression of a bacteriophage lysis gene, ▫X174 E or E-lysis gene.^24^ Later studies further validated that the strategy developed by Din et al. can be used to deliver nanobodies and other protein-based payloads.^25^ While the proposed strategy for protein secretion is a good choice for cancer related applications, as demonstrated in the mentioned studies, it may not be suitable for other applications, particularly those that require anti-inflammatory activity. Spontaneous lysis of GNB releases dsDNA and other intracellular aside from the payload protein(s) content which can cause an inflammatory response. In addition, payload protein is released to the environment with a sudden peak in concentration which can lead to local drug toxicity and prevent sustaining the concentration of payload at its therapeutical window. We proposed a controlled protein release system in GNB which relies on enzymatic cleavage of surface displayed proteins with a site-specific protease. The system has a titratable protein secretion profile; in other words, the amount of secreted protein can be tuned with a small molecule, e.g., aTc. Owing to its simplicity, the secretion system can be functionally implemented in complex genetic circuits.^8^ The main disadvantage of our system was the use of TEV protease, whose activity is affected by pH, temperature, and other environmental factors. Despite these efforts improved some drawbacks of native systems, proposed secretion systems still are limited to certain applications. Therefore, it is necessary to develop an efficient protein secretory production strategy that can easily be implemented in other GNB species.

Here, we present a novel synthetic secretory protein production strategy called iLOM-SS, which stands for **i**nducible **<u>L</u>**eaky **<u>O</u>**uter **<u>M</u>**embrane based **<u>S</u>**ecretion **<u>S</u>**ystem. In iLOM-SS, structural protein(s) at the periplasm are repressed by small RNAs to slightly destabilize the outer membrane integrity, thereby releasing proteins translocated at the periplasm to the extracellular milieu via free diffusion. To demonstrate and test the protein secretion with this strategy, we chose *E. coli* strains to construct the iLOM-SS by repressing the *lpp* gene (Figure 1a). We showed that proteins up to 100kDa can be secreted by iLOM-SS with detectable amounts minutes after the activation. Furthermore, protein secretion with the iLOM-SS can be titrated with a small molecule, and the protein secretion can be halted if the repression of *lpp* gene is relieved. We constructed a compact version of iLOM-SS and validated its functionality in different genetic circuit architectures. As a proof-of-concept, iLOM-SS was utilized for the secretion of a human cytokine, IL-1Ra from EcN strain. Considering the capabilities of existing secretion systems, the constructed iLOM-SS provides an efficient and safe way to secrete proteins in EcN. Moreover, the iLOM-SS can be constructed in other physiologically relevant GNB for protein secretion which can inaugurate the use of these microbes as shuttles to deliver of payload proteins at desired sites such as gut and tumors.

**Figure 1.**
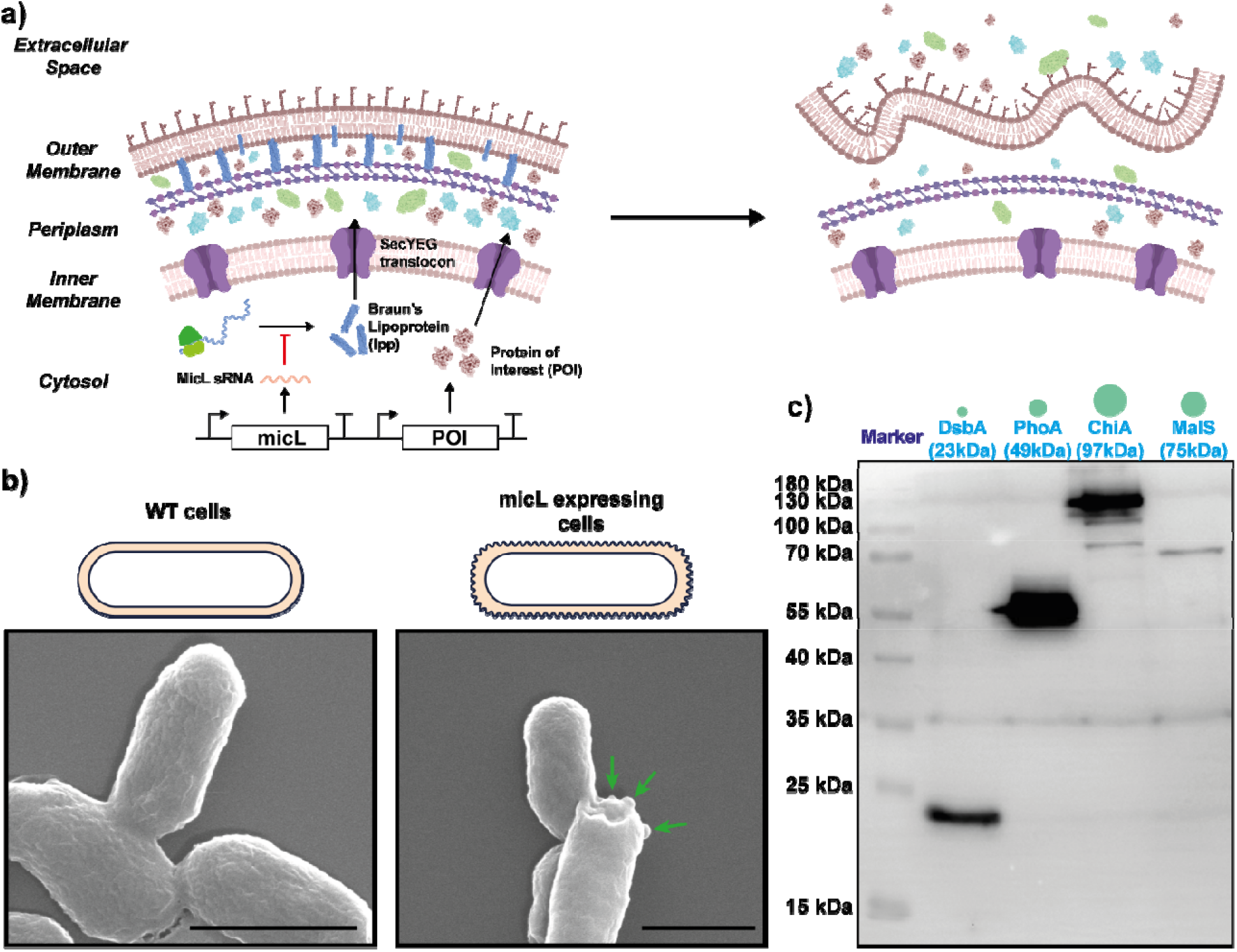
Development and validation of inducible leaky outer membrane (iLOM)-based secretion system in *E. coli*. a.) Schematic presentation of iLOM-based secretion system in *E. coli*. In the system, protein of interest (PoI) is translocated to the periplasmic space with either TAT or Sec-dependent secretion signal peptides (SP), e.g., pelB SP, dsbA SP, TorA SP. Leaky outer membrane was induced with expression of micL sRNA which exclusively inhibits the translation of Braun’s lipoprotein (lpp). Subsequently, it causes bleb formation on the outer membrane and leads to free diffusion of periplasmic content to the extracellular space. b) Morphology of *E. coli* cells under ESEM following the micL expression. The blebs were shown with green arrows on the micL expressing cells. Black bars are 1µm. c) Secretion of proteins, which have different molecular weights, with the iLOM-SS. The coding region of DsbA (23 kDa), PhoA (49 kDa), MalS (75 kDa) and ChiA (97 kDa) were cloned into expression vector with 6x his-tag. Following to the induction of micL sRNA and the proteins, supernatants were collected and directly loaded into SDS-PAGE gel with any pre-enrichment step. Anti-his antibody was used to collect Western Blot image.

## RESULTS

### Construction of the iLOM-SS in *E. coli*

To build the iLOM-SS, we manually searched the literature to find genes whose knock-out mutant causes the leaky outer membrane (LOM) phenotype in *E. coli* strains. Interestingly, the *lpp* gene that is shown as one of the mutated genes for LOM phenotype is exclusively repressed by a natural sRNA, named as micL.^26,27^ MicL sRNA was cloned into an IPTG inducible plasmid to test whether the expression of micL is sufficient to induce LOM phenotype in *E. coli* DH5α strain. Bacteria cells that express the micL sRNA and WT cells were grown overnight, and ESEM samples were prepared from grown bacteria cultures. Under ESEM, blebbing which was previously shown on *lpp* knock-out mutants and other *E. coli* mutants with LOM phenotype^28^ was observed on the outer membrane of micL expressing cells. (Figure 1b)

After validating the LOM phenotype induction with micL expression, outer membrane leakiness was analyzed with proteins that have different molecular weights. To do so, dsbA (23kDa), phoA (49kDa), malS (75kDa), and chiA (97kDa) genes were tagged with 6xHis tag and cloned separately into IPTG inducible micL expression plasmid under aTc inducible promoter. The plasmids were transformed into E. *coli* DH5α strain and cells were induced overnight with both aTc and IPTG to express micL and the proteins. The next day, cell supernatants were collected with centrifugation. Collected supernatants without any pre-enrichment methods, such as TCA precipitation, ultrafiltration etc. were analyzed with western blot using anti-6xHis tag antibodies. Western blot analysis confirmed that all four tested proteins were released from periplasm to extracellular space following to the micL expression. (Figure 1c). Furthermore, we also tested the free diffusion of FTIC conjugated 4kDa dextran polymer into the periplasm of micL expressing cells. Both microscopy and spectroscopy results revealed that micL expressing cells had more green fluorescence compared to WT cells thereby indicating that the FTIC conjugated polymer was retained in micL expressing cells. (Figure S1) These results suggested that micL expression induces LOM phenotype in *E. coli*.

### Characterization of the iLOM-SS

We characterized the iLOM-SS by secreting alkaline phosphatase (ALP) enzyme that is encoded in the phoA gene as ALP can degrade the colorless para-nitrophenol phosphate (PnPP) substrate to yellow colored para-nitrophenol (PnP) whose amount can be determined easily via measuring the absorbance of solution at 405nm. (Figure 2a) Firstly, the functionality of iLOM-SS was tested in different strains of *E. coli*. For this purpose, the reporter plasmid composed of aTc inducible phoA and IPTG inducible micL sRNA was transformed into different strains of *E. coli* including WT strain K-12, BL21 (DE3), and EcN. Cells were grown overnight and induced to express ALP and micL sRNA. Subsequent to cell induction, the ALP activity of cell supernatants was determined. For all strains, micL expression led to significantly higher ALP activity compared to uninduced cells. (Figure 2b) The results clearly indicate that the iLOM-SS is functional in different strains of *E. coli*. EcN strain was chosen as the model strain for further characterization experiments because of its probiotic activities.

**Figure 2.**
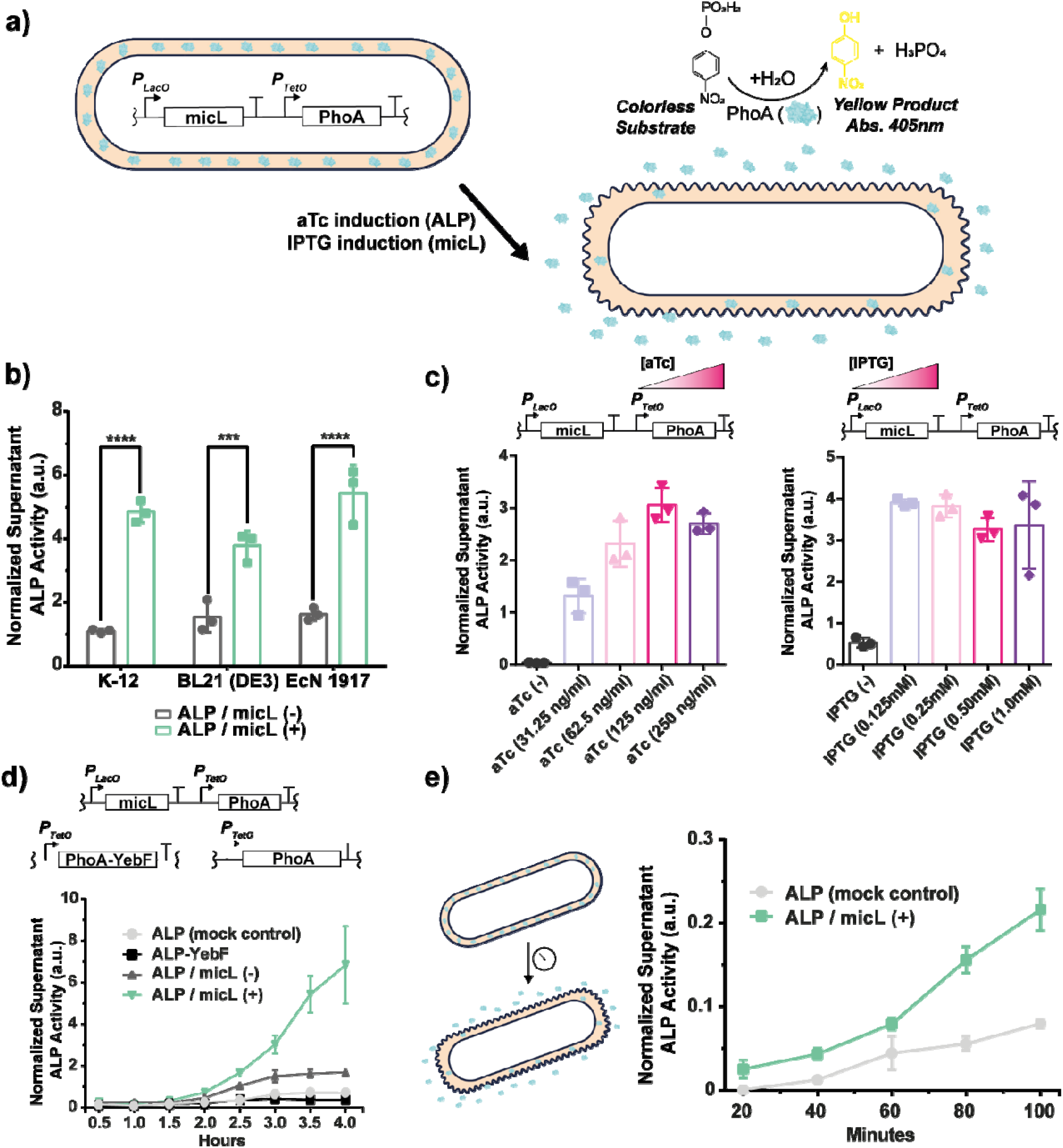
Characterization of iLOM-based secretion system in EcN. a) Schemati presentation of enzyme model used to detect secreted proteins amount in the extracellular milieu. PhoA encodes an alkaline phosphatase enzyme which degrades the colorless PnPP to yellow-colored nPP. Enzymatic activity of ALP was used to assess and compare secreted protein amounts in supernatants upon induction of micL and PhoA. b) Validation of iLOM-based secretion system in different strains of *E. coli*. The plasmid that encodes micL and PhoA wa transformed into *E. coli* K-12, BL21 (DE3) and EcN strains. Following induction with aTc and IPTG, ALP activity of supernatants obtained from cells was measured with PnPP degradation assay. Student’s t-test was used to assess the significance. c) Expression titration of micL with IPTG and PhoA with aTc to evaluate the controllability of iLOM-based secretion in terms of secreted protein amount. During titration experiments, the non-titrated inducer was added at maximum concentration which was 1mM for IPTG and 250ng/ml for aTc. Four different concentrations were determined for each inducer. d) Comparison iLOM-based secretion of ALP with YebF carrier protein-based secretion and non-secreting control in time domain. Supernatants of cells were collected in each 30 minutes for 4 hours to determine secreted ALP activity in the extracellular space. e) Determination of time required for activation of the iLOM-SS after the induction of micL. PhoA was induced beforehand to translocate ALP enzymes to the cell periplasm. Then, cells were diluted 100-fold into fresh media supplemented with both aTc and IPTG. Subsequently, supernatants were collected for 100 minutes with 20 minutes intervals. ALP activity of obtained supernatants was determined with PnPP assay. All experiments were conducted with three independent biological replicates.

Next, we tested whether the iLOM-SS could provide a titratable protein secretion when used in EcN. Bacterial cells carrying the reporter plasmid were induced by titrating either aTc or IPTG in four different concentrations meanwhile the other inducer that is not titrated was added at the maximum concentration which is 250ng/ml for aTc and 1mM for IPTG. After 4-hour induction, cell supernatants were collected to determine the ALP activity. The activity results revealed that titration of aTc thereby titration of phoA expression provided to control the amount of secreted ALP. On the other hand, titration of micL sRNA expression with IPTG did not provide the titratable protein secretion for the concentrations tried in the experiment. (Figure 2c)

We also benchmarked secretory capacity of the iLOM-SS against the widely used secretory fusion tag, YebF^29^, in *E. coli*. For comparison, phoA gene was inserted to C-terminal of YebF and only phoA expressing cells were also added in the experimental setup as a mock control. Overnight grown cultures of bacterial cells that carry the plasmids were diluted into fresh media and induced when their OD_600_ reached 0.1. After the induction, cell supernatants were collected in 30 minutes intervals for 4 hours to determine the ALP activity. Activity results showed that iLOM-SS when induced, could release a detectable amount of ALP enzyme to extracellular media as soon as 2 hours, and protein accumulation at the extracellular milieu continued throughout the experiment timeline. In contrast, YebF fusion tag failed to translocate the ALP enzyme same as the mock control. (Figure 2d)

Lastly, we determined the time required for the activation of iLOM-SS. Although a detectable amount of the ALP enzyme was found to be released at least 2 hours after the induction, we hypothesized that the production of ALP could be the bottleneck rather than the activation of iLOM-SS. To test this, bacterial cells were induced overnight to express ALP. The next day, cells were diluted into fresh media at the OD_600_ of 0.8, and micL sRNA expression was induced to activate the iLOM-SS. Subsequently, cell supernatants were collected every 20 minutes for 100 minutes to assess extracellular ALP amount with the activity assay. Accumulated ALP activity differs greatly in the supernatant of micL expressing cells compared to the mock control 60 minutes after the induction consequently indicating that iLOM-SS activation happens approximately one hour after the expression of micL sRNA. (Figure 2e)

### Stop switch design for the iLOM-SS

We built and validated a stop genetic switch that can halt the secretion with iLOM-SS in case of need. Our main hypothesis was that the LOM phenotype could be reversed therewith ceasing the secretion if lpp translation was rescued in the absence of micL sRNA expression. To test this, a toggle genetic switch that expresses micL sRNA with pTetO and mCherry with pLacO was constructed. In doing so, two states were defined with a toggle switch which are micL sRNA expressing secretory state and mCherry expressing non-secretory state. As secretion reporter, the plasmid that encodes arabinose inducible ALP was co-transformed into the toggle switch carrying cells. (Figure 3a) Cells were induced sequentially with aTc and IPTG for six days to change their states between secretory and non-secretory. After each induction, cell state change was confirmed by measuring the mCherry signal from the cells. As expected, cells were expressing mCherry following the induction of IPTG which changes the cell state to the non-secretory, and mCherry signal decreased when cells were induced with aTc which changes the cell state to the secretory. (Figure 3b and 3c) For further validation, cells in each state were induced with arabinose to check whether ALP could be released from the cells. Based on the ALP activity results, cells in the secretory state could release the ALP to extracellular. By contrast, no ALP activity was observed in the supernatant of cells whose state was the non-secretory. (Figure 3b) These results demonstrated that our hypothesis was correct, and an emergency genetic circuit can be constructed to stop the protein release from iLOM-SS by blocking the expression of micL sRNA.

**Figure 3.**
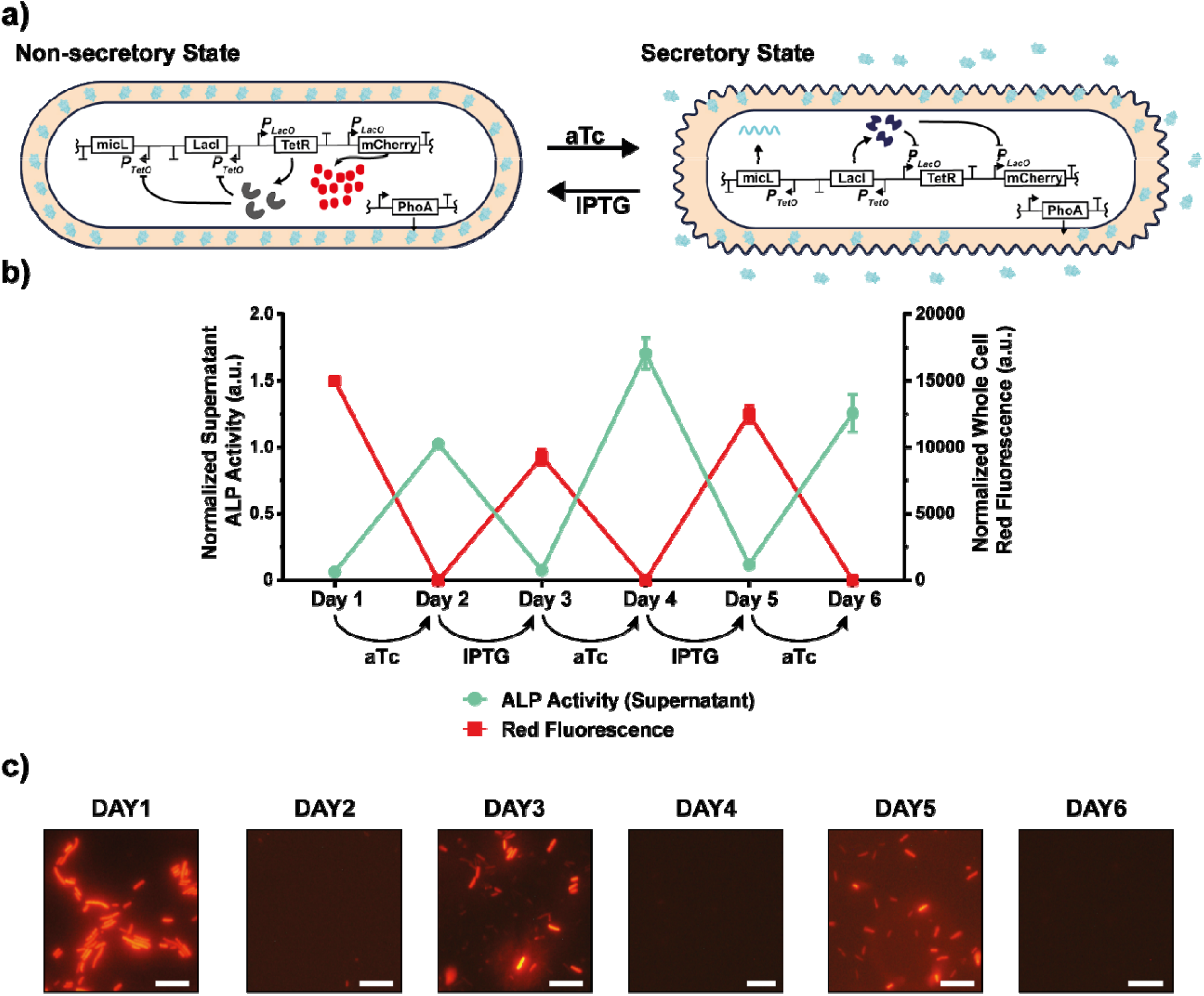
Leaky membrane can be restored in the absence of micL expression to halt the secretion of periplasmically translocated proteins with the iLOM-SS. a) Schematic representation of genetic circuit that was used to test the hypothesis of leaky membrane restoration. A toggle switch gene circuit with LacI/TetR repressor pair was constructed. The coding region of micL sRNA was placed downstream of pTetO promoter whereas mCherry expression was controlled by pLacO. In the non-secretory state, cells were expected to produce mCherry and TetR proteins; on the other hand, micL sRNA and LacI were expressed by the cells in the secretory state. b) Supernatant ALP activity and whole cell red fluorescence of the cells sequentially induced by IPTG and aTc for six days. c) Red fluorescence microscopy images of the cells each day. White bars are 5µm. All experiments were conducted with three biological replicates.

### Development of the compact iLOM-SS and its functionality in different genetic circuit architectures

To decrease the metabolic load on cells, we engineered a compact version of iLOM-SS where expression of POI and micL sRNA takes place on the same transcriptional unit thereby eliminating the necessity of second expression cassette. To do so, micL sRNA was flanked by two self-cleaving ribozymes, hammerhead ribozyme and hepatitis virus ribozyme^30^, and resulting RNA named as c-micL was directly fused to the 3’ of mRNA region coding the POI to construct a single transcription unit (STU) for compact iLOM-SS. (Figure 4a) PhoA was utilized to test the functionality of the compact version of iLOM-SS. Supernatant ALP activities demonstrated that cell expressing the phoA fused with c-micL can release ALP enzyme to extracellular media significantly higher than the mock control. (Figure 4b)

**Figure 4.**
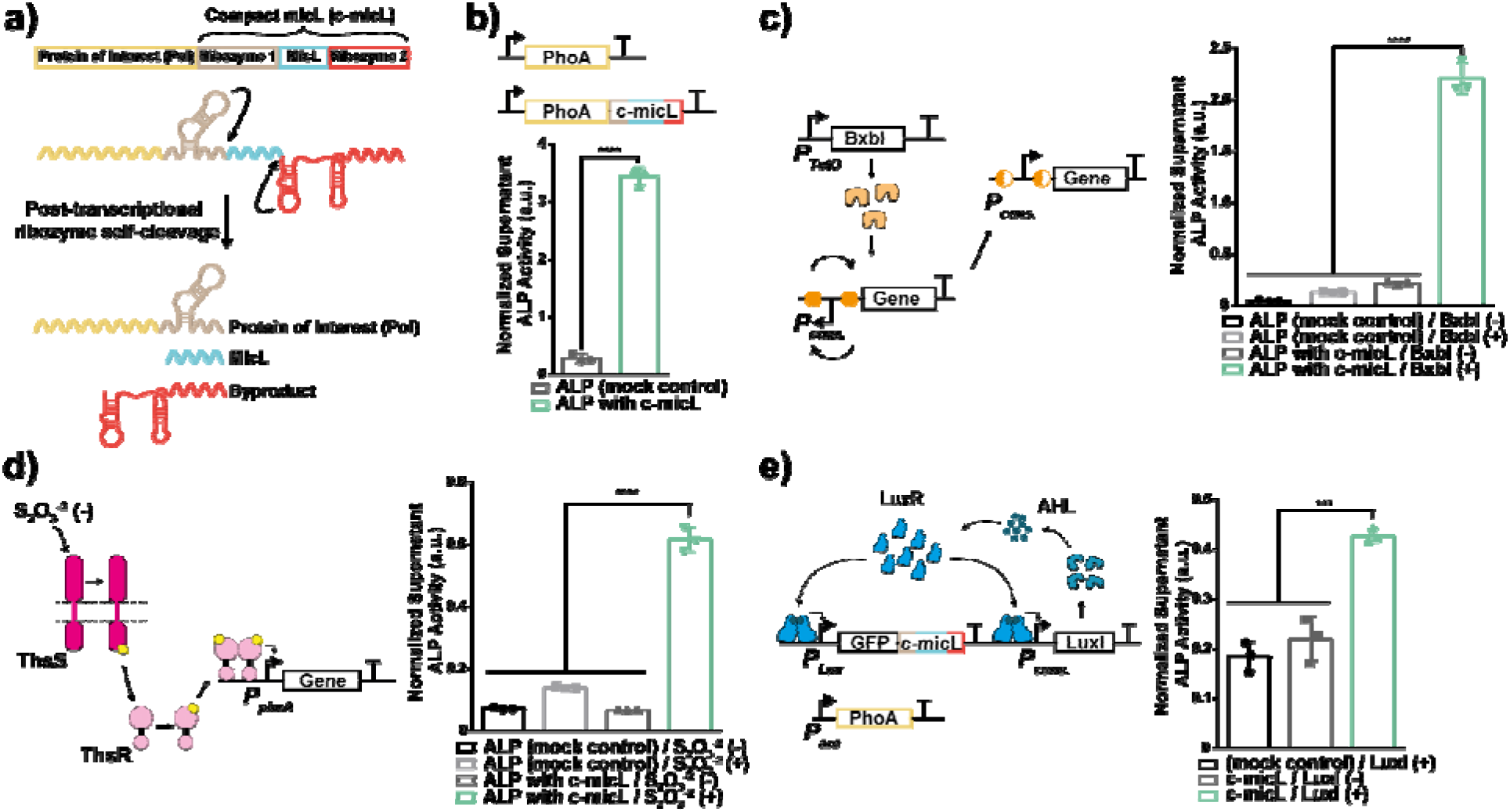
Development of single transcription unit for iLOM-based secretion, and its functional validation in different genetic circuit architectures. a) Visual depiction of single transcription unit topology for the iLOM-SS. Coding region of micL sRNA is flanked by hammerhead ribozyme (ribozyme 1) at its 5’ end and hepatitis delta virus (HDV) ribozyme (ribozyme 2) at its 3’ end. The constructed DNA region is named compact micL (c-micL) and directly fused to 3’ end of PoI coding mRNA. After the transcription, ribozyme 1 and ribozyme 2 catalyze the cleavage of phosphodiester bonds at their 3’ and 5’ ends, respectively. The self-processing of ribozymes results in formation three RNA molecules; PoI coding mRNA fused with ribozyme 1, micL sRNA and ribozyme 2. b) Functional validation of iLOM-based secretion of ALP with c-micL. Supernatant activities of cells that were carrying either only PhoA gene (mock control) or PhoA fused with c-micL were measured. c, d, e) Performance and activity validation of iLOM-based secretion in recombinase-based memory circuit (c), two-component system that senses presence of thiosulfate (d), and pLux/LuxI quorum sensing circuit from (e). For (c) and (d) Either only PhoA gene (mock control) or PhoA fused with c-micL was inserted at the downstream of the promoters. For (e), c-micL was fused to the GFP coding sequence and PhoA expressed was controlled with arabinose inducible promoter. All experiments were carried out with three biological replicates.

We next explored the functionality of compact iLOM-SS design in different genetic circuit architectures that can exploited for variety functions such as memory recording, crowd, and small molecule sensing. To begin with, ALP secreting compact iLOM-SS design was implemented in recombinase-based genetic circuit where expression of a gene is driven by recombinase-assisted inversion of inverted promoter. (Figure 4c) Secondly, the design was utilized in two component system (TCS) found in bacteria which senses extracellular clues through membrane spanning sensor protein and transfer this information its cognate intracellular transcription factor to induce gene expression. (Figure 4d) Lastly, quorum sensing circuit from *Vibrio fischeri* was used to test the compact iLOM-SS. (Figure 4e) For all circuit architectures mentioned above, compact iLOM-SS carrying cells released significantly higher ALP enzyme to extracellular media compared to control groups. (Figure 4c, 4d, 4e) Taken together, these results suggest that compact iLOM-SS is a valid strategy to secrete proteins and can be implemented in complex genetic circuits where protein secretion is required as an output.

### Secretion of Bioactive Human Cytokine, IL-1Ra, with iLOM-SS

As a proof-of-concept, we sought to secrete a therapeutic protein using the iLOM-SS with EcN. Human cytokine interleukin 1 receptor antagonist (IL-1Ra) was chosen as a model protein since it is an FDA-approved drug used in rheumatoid arthritis and other inflammatory diseases, and there are well-established analytical tests to determine its amount and biological activity. Secretion plasmid was constructed by changing the phoA gene with IL-1Ra coding region with pelB periplasm targeting signal in micL/phoA expression plasmid. Secretion of IL-1Ra by bacterial cells was determined with both ELISA and western blot, and biological activity of IL-1Ra was measured using the A375.S2 cell model in which IL-1α induced apoptosis was inhibited.^31^ (Figure 5a) Both ELISA and western blot results validated that IL-1Ra was secreted to extracellular media with iLOM-SS; furthermore, at least a 7-fold increase in the amount of extracellular accumulated IL-1Ra was observed compared to control group that only expresses IL-1Ra at its periplasm. (Figure 5b and 5c) When added onto IL-1α treated A375.S2 cells, secreted IL-1Ra from EcN can prevent cell death with dose-dependent manner. (Figure 5d) These results highlighted that iLOM-SS can secrete pharmaceutically active proteins.

**Figure 5.**
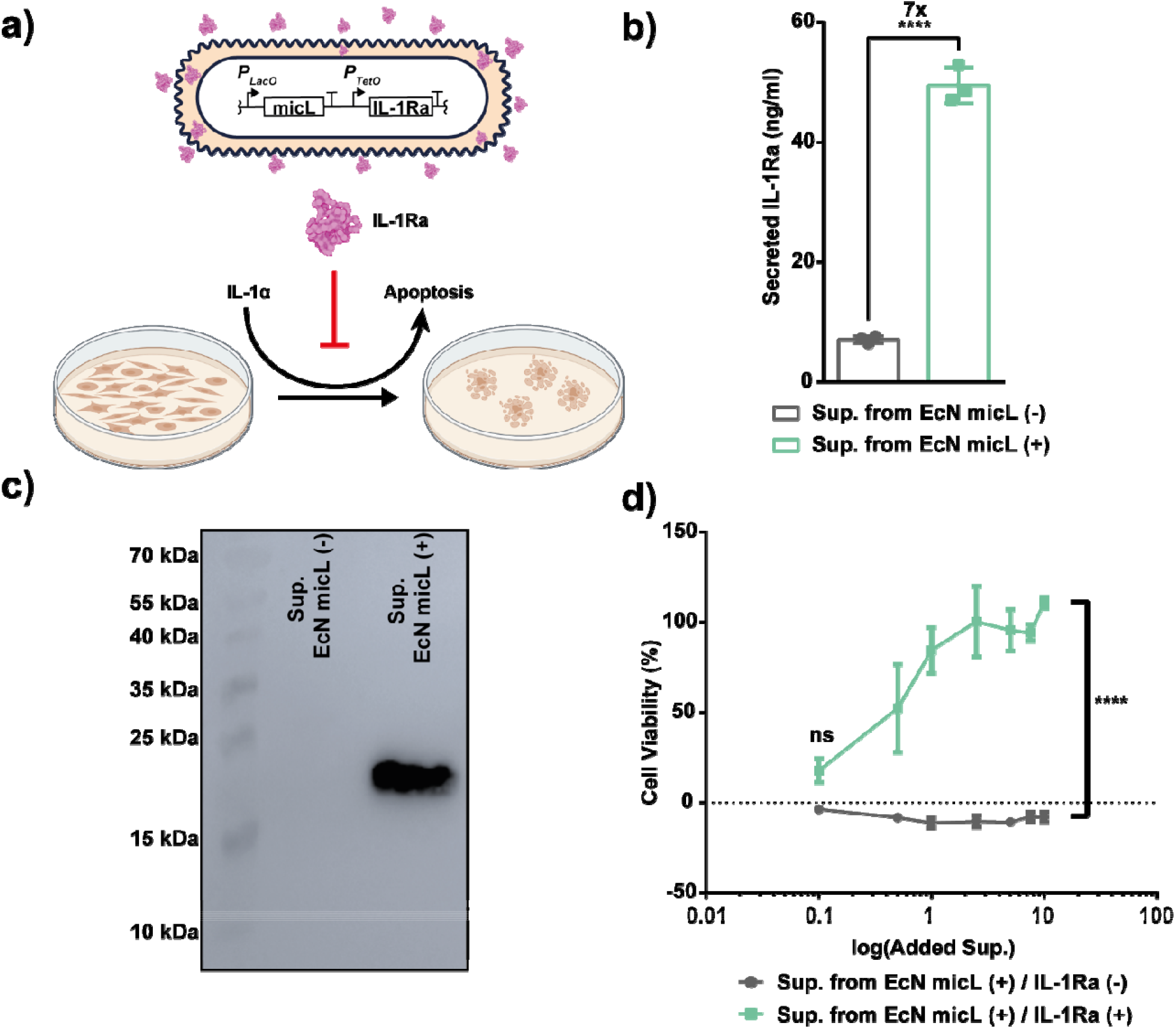
Active IL-1Ra secretion with iLOM-based secretion system using EcN. a) Schematic presentation of the genetic circuit to secrete IL-1Ra with iLOM-based secretion system and *in vitro* assay to determine the activity of IL-1Ra. The activity of secreted IL-1Ra was determined using A375.S1 cells. In the presence of active IL-1Ra, IL-1α induced apoptosis of A375.S2 cells are inhibited which can be detected by measuring cell viability after the IL-1α treatment. b) Secreted IL-1Ra amount to extracellular space with iLOM-based secretion system. The concentration was determined via IL-1Ra specific ELISA without any enrichment method e.g., ultrafiltration. c) Verification of IL-1Ra at the supernatant following the induction. Anti-IL-1Ra antibody was used to capture the Western blot image. d) Cell viability of A375.S2 cells determined by utilizing MTT assay after the treatment with IL-1α and the supernatant obtained from either IL-1Ra secreting EcN cells or control cells. The experiments were conducted with at least three independent replicates.

In addition, we tested whether the leaked periplasmic content (LPC) from EcN during the secretion of the cargo protein causes any toxicity *in vitro* on the Caco-2 cell line. We proved that LPC has no cytotoxic and pro-inflammatory effects and didn’t affect the migration of Caco-2 and permeability of the Caco-2 monolayer. (Figure S2 and Figure S3) Also, we showed that expression of micL didn’t cause growth defects on the EcN strain. (Figure S4a) Surprisingly, the invasion of Caco-2 cells by EcN as well as cell motility of EcN decreased following the micL expression. Based on the data obtained with Caco-2 cells, we claimed that iLOM-SS is a safe strategy to engineer EcN cells as sentinels to deliver protein therapeutics in the gut.

## DISCUSSION

Here, we engineered a protein secretion system which didn’t exist in nature for *E. coli* strains to release wide range of proteins into extracellular space. In the secretion system, proteins that are translocated to periplasm are released into the extracellular milieu by induction of LOM phenotype through translational suppression of *lpp* protein by micL sRNA. We envisioned this system to overcome the drawbacks of natural secretion systems such as high metabolic load and narrow substrate profile. We demonstrated that the secretion system called iLOM-SS is functional in other strains of *E. coli* including industrially renowned BL21 (DE3) protein expression strain^32^, and clinically proven probiotic EcN strain.^33^ Although we focused characterizing the iLOM-SS in EcN strain for therapeutic cargo delivery, the versatility of iLOM-SS among *E. coli* strains can expand its applicability in different areas where protein secretion is required.

Titratable action and stop switches are desirable features for complex biological therapy agents, especially for living therapeutics, because life-threating adverse reactions some of which cannot be predicted prior to human clinical trials might occur after administration.^34^ Therefore, there are substantial efforts to design genetic switches for controlling the action of advanced biological agents such as engineered bacteria, mRNA and CAR-T cells.^35–38^ We showed that the secreted protein amount with iLOM-SS is titratable with a small molecule inducer. Although the titration experiments were conducted with aTc which is not applicable for human administration, there are promoter/transcriptional factor pairs whose inducers are safe to use in humans.^39^ We also demonstrated that a stop switch can be used to halt the protein secretion with iLOM-SS in case of need. Altogether with these tools, it is possible to control the secretory action of iLOM-SS augmented microbes with external clues however, future studies are required to confirm their performance *in vivo*.

Many physiological processes in the human body such as blood sugar regulation necessitate genetic tools with fast response time, yet current tools whose output are linked with transcription and translation of a gene can’t provide the required responsiveness.^40^ We showed that iLOM-SS can release proteins within minutes following the activation if the released protein is translocated to periplasm beforehand. Thus, our secretion system encourages designing bacteria as protein delivery systems for applications that need fast response.

The modularity of genetic tools is one of the core design concepts in biological engineering as it accelerates the design-build-test cycle by enabling the construction of different genetic circuits spontaneously.^41^ For modular implementation, we developed a compact version of the iLOM-SS where micL sRNA can be expressed within the same transcriptional unit as the protein-of-interest. To demonstrate the modularity of compact iLOM-SS, its functionality was validated in different genetic circuit architectures which can sense small molecules or record the presence of a small molecule inducer. We believe that compact iLOM-SS will provide a platform for researchers and synthetic biology enthusiasts to design genetic networks with protein secretion as an output. Owing to the simplicity of the compact iLOM-SS with plug-play design, the democratization of protein secretion will further stimulate synthetic biology research in which protein secretion is a prerequisite.

Although, as a proof of concept, we showed the secretion of IL-1Ra in this study, we believe that iLOM-SS augmented EcN can be used to develop sentinel bacteria for treatment of variety gut pathologies such as inflammatory bowel disease, celiac disease, etc. There is a vast amount of literature present showing that *E. coli* strains can express a variety of proteins recombinantly in their active forms.^42–46^ In addition, we demonstrated that leaked periplasmic content of EcN during secretion with the iLOM-SS has no pro-inflammatory or cytotoxic effects *in vitro* on Caco-2 human epithelial cell model thereby indicating that LPC doesn’t have any apparent negative impacts on human cells and its presence can be tolerated. However, these findings should be further validated *in vivo* models with additional parameters to ensure the safety of protein delivery with iLOM-SS augmented EcN.

We anticipate that the general approach that is used for the development of iLOM-SS can be exploited to engineer non-model GNB strains for protein secretion. There are emerging strains such as Pseudomonas putida^47^, Komagataeibacter spp.^48^, and Bacteroides spp.^49^ can be enhanced with iLOM-SS for bioremediation, biomaterials production, and therapeutical applications, respectively. One can design an iLOM-SS for non-model GNB through a simple CRISPRi-based screening.^50^ With the vast amount of literature and readily available tools, we forecast that developing iLOM-SS in other GNB should take a 6-month time frame.

To conclude, we presented an approach called iLOM-SS, to develop a synthetic secretion system for GNB to mitigate the problems of naturally evolved secretion systems. Based on this approach, a secretion system was constructed and characterized for *E. coli* strains. We showed that the constructed secretion system offers many features such as modularity, controllability, and versatility. We expect that iLOM-SS will ease the burden on designing GNB for many applications such as live shuttles for therapeutical protein delivery.

## METHODS

### Plasmid Construction, and Strains

All plasmids used in this study were constructed with Gibson Assembly. Plasmid sequences were verified with either Sanger sequencing or NGS. All enzymes used in this for plasmid construction were purchased from New England Biolabs (NEB). Primers were ordered from Oligomer Inc. and BiOligo Inc. If needed, gene synthesis service of Genewiz Inc was utilized. All plasmid maps, primers, and sequencing results were given in supplementary information. LB media and LB agar with appropriate antibiotics were used for bacteria cultivation unless otherwise noted. DH5α with Pro cassette was used for experiments in Fig1b, Fig1c, and Fig4. BW25113 ΔLacI was used for experiments in Fig3. For all other experiments, EcN 1917 strain was used. Bacteria cells were grown in a shaking incubator with liquid media if not otherwise indicated.

### Scanning Electron Microscopy

Bacteria cells bearing the plasmid that expresses micL sRNA under pLacO promoter were induced overnight in the presence of 1mM IPTG at 37 °C in LB liquid media with shaking. After the induction, cells were collected by centrifugation and washed two times with PBS. Cell pellet was resuspended in PBS at 10^9^/ml concentration. A small piece of isopropanol washed silica wafer was placed at the bottom of the 96 well plate and 10µl of PBS suspended bacteria was put on the silica wafer. The droplet was incubated for 10 minutes; 200µl of 2.5% glutaraldehyde (diluted in PBS) solution subsequently was added into well. Samples were fixed in glutaraldehyde solution on top of silica plate overnight at 4 °C. Next morning, the silica plate was washed 3 times with PBS followed by 3 times with ddH_2_O and once with ethanol solutions (25%, 50%, 75%) for 5 minutes each. Lastly, silica plate was washed three times pure ethanol and dried with critical point dryer. After drying, wafer piece with bacteria was mounted on the SEM stub and coated with Au at 8nm thickness. For visualization, Environmental SEM (Tecnia) was utilized.

### Western Blot Analysis for DsbA, ALP, MalS, and ChiA Secretion with iLOM-SS

Cells carrying the plasmids for N-terminal histidine tagged recombinant versions of DsbA, ALP, MalS and ChiA was induced with 1mM IPTG to express micL sRNA and 250ng/µl aTc to express the proteins for overnight at 37°C in shaking incubator. Following the induction, cells were centrifuged at maximum speed for 5 minutes to collect cell-free supernatant. 20µl of cell supernatant for each protein was mixed with Laemmli SDS-PAGE loading dye and loaded into wells of SDS-PAGE (12%) after boiling for at minutes at 95 °C. Proteins were transferred to nitrocellulose membrane with Bio-Rad’s Trans-Turbo Blot System using the Turbo-Blot RTA kit. Anti-histidine mouse IgG at 1:15000 dilution and anti-mouse IgG goat IgG conjugated with HRP at 1:10000 were utilized to obtain the western blot image. Vilber Gel Imaging System was used for imaging.

### Determination of Supernatant ALP Activity

Following the induction of cells, cell-free supernatant was collected as described above. 50µl of cell-free supernatant was mixed with 200µl of 5mM PnPP solution (PnPP dissolved in 1 mm MgCl2, 1 mm ZnCl2, 0.1▫m glycine, pH 10.4). The solution was incubated at 37 °C for 1 hour. Absorption at 405nm was measured to assess the formed PnP. To normalize the data to cell number, absorption values obtained from the activity assay were divided by cell optical density at 600nm. The M5 spectrophotometer (Molecular Devices) was used for all measurements.

### Characterization of iLOM-SS with ALP

To characterize the iLOM-SS in different *E. coli* strains, bacterial cells were transformed with the plasmid that encodes IPTG inducible micL and aTc inducible phoA. Single colony transformants on selective agar were picked and grown overnight in the presence of appropriate antibiotics. The next day, cells were rediluted 1:100 into fresh media, grown until the OD_600_ of bacteria culture reaches 0.1, and induced 1mM IPTG and 250ng/µl aTc for 4 hours. After the induction, cell supernatants were collected with centrifugation at maximum speed for 5 minutes. ALP activity of cell supernatants was determined as described above.

In inducer titration experiments, EcN cells carrying the phoA/micL expression plasmid were grown the same as above. At the induction step, the concentration of one of the inducers was kept at the maximum induction concentration which was 1mM for IPTG and 250ng/µl for aTc, meanwhile, concentration of the other inducer was titrated. The titration range for IPTG was 0.125mM, 0.25mM, 0.5mM, and 1mM; on the other hand, the range for aTc was 31.25ng/µl, 62.5ng/µl, 125ng/µl, and 250ng/µl. Following the induction, ALP activity of cell supernatants was determined same as above.

For comparison of iLOM-SS with YebF and the mock control experiments, phoA/micL expression plasmid, phoA expression plasmid without micL, and only phoA expression plasmid were transformed into EcN strain separately. Cells were grown and induced with IPTG and aTc their maximum induction concentration. Following the induction, Cell supernatants were collected in each 30 minutes for 4 hours. ALP activity of collected cell supernatants was determined.

To find the time required for iLOM-SS activation, EcN cells that were transformed with phoA/micL expression plasmid was grown overnight in presence of 250ng/µl aTc. Cells were rediluted at OD_600_ of 0.8 into fresh media supplemented with antibiotics and both of the inducers. After the dilution, cell supernatants were collected in every 20 minutes for 100 minutes and ALP activity of supernatants was determined.

### Consecutive Inductions of the iLOM-SS in Toggle Switch

Previously deposited pECJ3 plasmid was used to construct the toggle switch used in the study.^51^ Cells were co-transformed with the toggle switch plasmid that encodes mCherry and micL sRNA, and the plasmid that encodes arabinose inducible ALP. Transformed cells were selected on LB agar plate supplemented with antibiotics. A single colony was picked and was grown overnight in M9 minimal media supplemented with 0.2% glycerol, 0.05% casamino acids, and antibiotics at 37°C in the shaking incubator. Cells in grown culture were diluted 1:10000 into fresh media supplemented with antibiotics and 1mM IPTG. Cells were induced for 24 hours at 37°C in the shaking incubator. The next day, cells were diluted into two fresh media in different tubes with 1:10000 and 1:100 dilution factors. 1:10000 diluted cells were induced with 250ng/µl aTc in the same condition as IPTG to change the state of the cells. Meanwhile, arabinose was added into the tube that contained 1:100 diluted cells to express ALP. Arabinose induction was carried out at 37°C for 4 hours in the shaking incubator. Following the arabinose induction, the ALP activity of the cell-free supernatant was determined as described above. In addition to that, whole cell red fluorescence was measured, and cells were visualized under fluorescence microscopy. The cell state was changed six times with consecutive induction of cells with IPTG and aTc. In each state change was validated with measuring the ALP activity of cell-free supernatants, and the whole cell red fluorescence, and visualizing cell under fluorescence microscope. Zeiss fluorescence microscope and M5 spectrophotometer (Molecular Devices) were utilized.

### Characterization of the compact iLOM-SS in Different Genetic Circuits

To validate the functionality of the compact iLOM-SS, plasmids that encode pTetO controlled either PhoA with c-micL or mock control PhoA were transformed into the strain. Single colonies were picked from agar plate and induced with 250ng/µl aTc for overnight in shaking incubator at 37 °C. The next day, supernatant ALP activities of cells were determined same as described above.

For functionality assay in recombinase-based memory circuits, cells were co-transformed with the plasmid that encodes aTc inducible BxbI recombinase enzyme^52^, and the plasmid that encodes either ALP with c-micL or ALP at the downstream of inverted proD promoter flanked by anti-parallelly aligned attP and attB sites of BxbI. Single colonies were picked agar plate and grown in LB liquid media with 250ng/µl aTc overnight at 37°C in the shaking incubator to express BxbI. Subsequent to overnight induction of BxbI, cells were diluted into fresh media at the 1:10000 dilution factor. Diluted cells were grown for 24 hours in liquid LB media supplemented only with antibiotics. The next day, ALP activity of cell-free supernatant of grown cells was measured same as described above.

To test the functionality of the compact iLOM-SS in thiosulfate sensing TCS sensor, cells were co-transformed with pKD236 and modified pKD237^53^ that expresses either ALP or ALP with c-micL under control of phsA promoter. Single colonies were picked from the agar plate and induced with 5mM sodium thiosulfate in liquid LB media overnight at 37 °C. ALP activity of cell-free supernatants obtained from induced and uninduced cells was determined described as above.

For functional validation of the compact iLOM-SS in LuxI/LuxR quorum sensing circuit^54^, cells carrying the plasmids that encode arabinose inducible PhoA, AHL inducible GFP fused with c-micL and LuxI with constitutively expressed promoter was grown for overnight in liquid LB media at 37 °C. The next day, cells were diluted with 1:100 dilution factor into a 1:1 mixture of fresh media and spent media that obtained from overnight culture. Arabinose is added into the media to induce ALP expression at 0.2% final concentration. The next day, ALP activity of cell-free supernatant of grown cells were measured same as described above. As controls, plasmids that either express GFP without c-micL or lack of LuxI expression cassette were used.

### IL-1Ra ELISA, Western Blot Analysis and Activity Measurements

Cells bearing the expression cassette for IL-1Ra and micL sRNA were induced with aTc and IPTG for overnight at 37 °C. Next day, cell-free supernatant was collected same as described above. ELISA (R&D Systems) was performed to determine secreted IL-1RA in the control and sample supernatants without any enrichment method e.g., ultrafiltration. 50x fold concentrated supernatants were used in western blot analysis and activity assays. Western Blot was performed same as described above except the IL-1Ra specific antibody was used instead of the anti-his tag antibody.

To validate the activity of secreted IL-1Ra, cell-free supernatant that contains secreted IL-1Ra was obtained same as described above. The supernatant was further concentrated by 50x fold with ultrafiltration. The desalting column was used to exchange the buffer of supernatant to PBS. The obtained solution was used in activity assays. A375.S2 cell line was used to determine the activity. 2000 cells per well were seeded into 96-well plate. Cells were incubated for a day prior to the treatment. On the day of treatment, 5ng/ml IL-1α was added to each well. Concentrated supernatant was added into wells in different amounts ranging from 0.1µl to 10µl. Following the three-day incubation, MTT assay was conducted to measure cell viability. Pure IL-1Ra, the supernatant obtained from micL sRNA expressing EcN and PBS were included in the assay as controls.

### Statistical Analysis

GraphPad Prism was used to visualize data and calculate p-values. 0.05 < p-value was taken as significant for all data obtained in the study. To compare values, appropriate statistical test was conducted such as Student’s t-test, one-way ANOVA, and two-way ANOVA. Each measurement was taken independently. At least, three replicates were used for all experiments. Symbols of ^*, * *, * * *^, and ^* * * *^ were used for p values that were smaller than 0.05, 0.01, 0.001 and 0.0001, respectively.

## Supporting information

Supplementaal Experimental Data

## DATA AVAILABILITY

## ACKNOWLEDGEMENTS

REA was supported by Bilkent UNAM Graduate Fellowship.

## AUTHOR INFORMATION

### Contributions

R.E.A. and U.O.S.S. conceived and designed the study. R.E.A., C.E.O and I.N.C. cloned the vectors and performed the experiments. R.E.A. analyzed the data under the supervision of U.O.S.S., and generated the figures. R.E.A. wrote the main draft. All authors reviewed and edited to finalize the manuscript. All authors approved the final version. U.O.S.S. acquired fundings for the study.

## ETHICS DECLARATIONS

UOSS and REA filed a patent application of tools and approaches developed in this study.

