## Supplementaal Experimental Data for "A Synthetic Protein Secretion System for Living Bacterial Therapeutics"

**SUPPLEMENTARY INFORMATION**

**SUPPLEMENTARY METHODS**

**FTIC-4kDa Dextran Diffusion Assay**

Cells were induced with IPTG for expression of micL sRNA, same as described in the main text. Following the induction, cells were collected and centrifuged at maximum speed. Cell supernatants were discarded; obtained pellets were washed twice with PBS to remove any residues of LB. Then, cells were resuspended in 10mg/ml FTIC-4kDa Dextran dissolved in PBS. Suspended cells were incubated at room temperature for 2 hours with shaking. After the incubation, FTIC-4kDa Dextran solution was removed via centrifugation. Cell pellets. were washed two with PBS after the treatment. Washed pellets were resuspended in PBS to measure whole cell green fluorescence and to visualize under a fluorescence microscope.

**Determination of Effects of Leaked Periplasmic Content (LPC) on Caco-2 Cell Viability, Monolayer Permeability, and Cell Migration**

For mammalian culture experiments, Caco-2 cells were expanded in high glucose DMEM supplemented with 10% FBS and 4mM L-Glutamine. Cells were grown in humified incubator with 5% CO_2_ at 37 ˚C for all experiments.

Caco-2 cells were treated with LPC containing supplemented DMEM to test safety parameters. To obtain the LPC-DMEM, EcN cells bearing the micL expression plasmid were diluted 1:100 into fresh DMEM and grown for 37 ˚C in shaking incubator for 3 hours in the presence of 1mM IPTG. Cells were centrifuged at full speed for 5 minutes at 4 ˚C to obtain the cell-free LPC-DMEM. To remove remaining cells after centrifugation, the media was filtered with 0.2µm filter. Subsequently, 10% FBS, and 4mM L-glutamine were supplemented to LPC-DMEM to obtain full growth media for Caco-2. To ensure the sterility, media was filtered again with 0.2µm filter. As a control, mock DMEM was obtained with similarly by using WT EcN.

For cytotoxicity assay, 100µl of suspended Caco-2 cells were seeded into 96 well plate at cell concentration of 10^5^ per well. Seeded cells were incubated for 24 hours before the LPC treatment. 100µl of supplemented LPC-DMEM was added into well on the day of experiment. As negative controls, mock DMEM control and untreated DMEM were used. Cells were treated for 3 days with LPC. At the end of experiments, MTT assay was conducted to compare the viable cell numbers in the treatment and control groups.

For permeability testing, Caco-2 monolayers were formed onto cell culture insert (Greiner-Bio, 662640). To do so, 10^6^ Caco-2 cells that were dissolved in 250µl growth media per insert were seeded onto apical side of inserts. Inserts were placed into 24-well plate and 750µl of growth media was added into basolateral side. Cells were grown for 21 days to induce spontaneous differentiation for monolayer formation. During the growth, media in both apical and basolateral sides were changed in two-day intervals. At the day 21, transepithelial electrical resistance was measured using Minicell ERS-2 Voltohmmeter (Millipore) complying with manufacturer’s instructions. Monolayers that had TEER value over 250 Ω.cm^2^ were selected for LPC treatment. Growth media at apical side was changed to mixture 1:1 of supplemented DMEM and LPC-DMEM while standard growth media was used for basolateral side. For control groups, normal growth media or DMEM obtained from WT EcN was used instead of LPC-DMEM. Cells were treated three days with LPC. TEER values for all groups were determined in each day during the treatment. At the end of the treatment, FTIC-Dextran diffusion assay was conducted with the monolayers. Growth media at the apical and basolateral sides were collected for IL-8 ELISA. 250µl and 750µl of HBSS buffer supplement with 1% glucose was added into apical and basolateral sides, respectively. 4kDa-dextran conjugated with FTIC was added to apical side at final concentration of 10µg/ml. Cells were incubated for 1 hour in the incubator. Following to the incubation, 200µl of solution from both sides were collected; fluorescence intensity of solutions were measured with spectrophotometer. As a diffusion positive control, empty cell insert was used.

For migration assay, Caco-2 cells were seed into 96 well plate and grown till they reached the full confluency. A small hole was created with pipette tip attached to vacuum after discarding the spent media. 100µl of fresh growth media was added into wells. Cells were incubated for 24 hours to recover at the hole edges. Subsequently, 100µl of LPC-DMEM was added into wells for treatment. Same controls used in previous experiments were also included in the migration assay. Cells were incubated for 3 days to observe the cell migration. In each day, picture of the holes was taken with a light microscope (Zeiss). Migration was calculated based on the closure percentage of hole compared initial hole dimensions. ImageJ software was used for calculations.

**Determination of Inflammatory Response of Treated Caco-2 Cells with LPC**

IL-8 secretion was determined from LPC treated differentiated and undifferentiated Caco-2 cells. Collected growth media from the differentiated cells used for permeability was used to determine secreted IL-8. For undifferentiated cells, 100µl of Caco-2 cell suspension were seeded into 96-well plate at 10^5^ cells per well seeding density and incubated for 24 hours for proper attachment. LPC treatment was conducted same as described in cytotoxicity assay. At the end of incubation, cell media was collected. IL-8 specific ELISA (R&D Systems) was conducted to determine secreted IL-8 amount following the LPC treatment.

**Determination of Growth Curve and Cell Motility**

To determine the growth curve, cells picked from single colonies on an agar plate were grown overnight. The next day, cells were rediluted into fresh media with 1:100 dilution factor and induced with 1mM IPTG. Optical density of cells at 600nm was measured every 20 minutes for 4 hours.

Cell motility was determined using soft agar method. LB agar plates that contain 0.3% agar were supplemented with 1mM IPTG and proper antibiotics if needed. 10µl of overnight grown culture was dropped at the center of plate. Plates were incubated at 37 ˚C for overnight. Plate images were taken with Vilber Imaging System.

**Caco-2 Invasion Assay**

Caco-2 cells were grown until the full confluency, same as described in the main text. At the day of the experiment, the spent growth media was discarded, and cells were washed 2 times with PBS gently. 200µl of a bacterial cell suspended in PBS whose an OD_600_ of 0.5 was added onto Caco-2 cells; then they were incubated at 37 ˚C for 4 hours. After the incubation, the bacterial cells were discarded, and Caco-2 cells were washed twice with PBS. 100µl of 0.2% Tween-20 dissolved in PBS was added onto cells to collect intracellular bacteria. Following an one hour, incubation, solutions were collected, serially diluted, and plated on agar plates to calculate CFU for each well.

**SUPPLEMENTARY FIGURES**

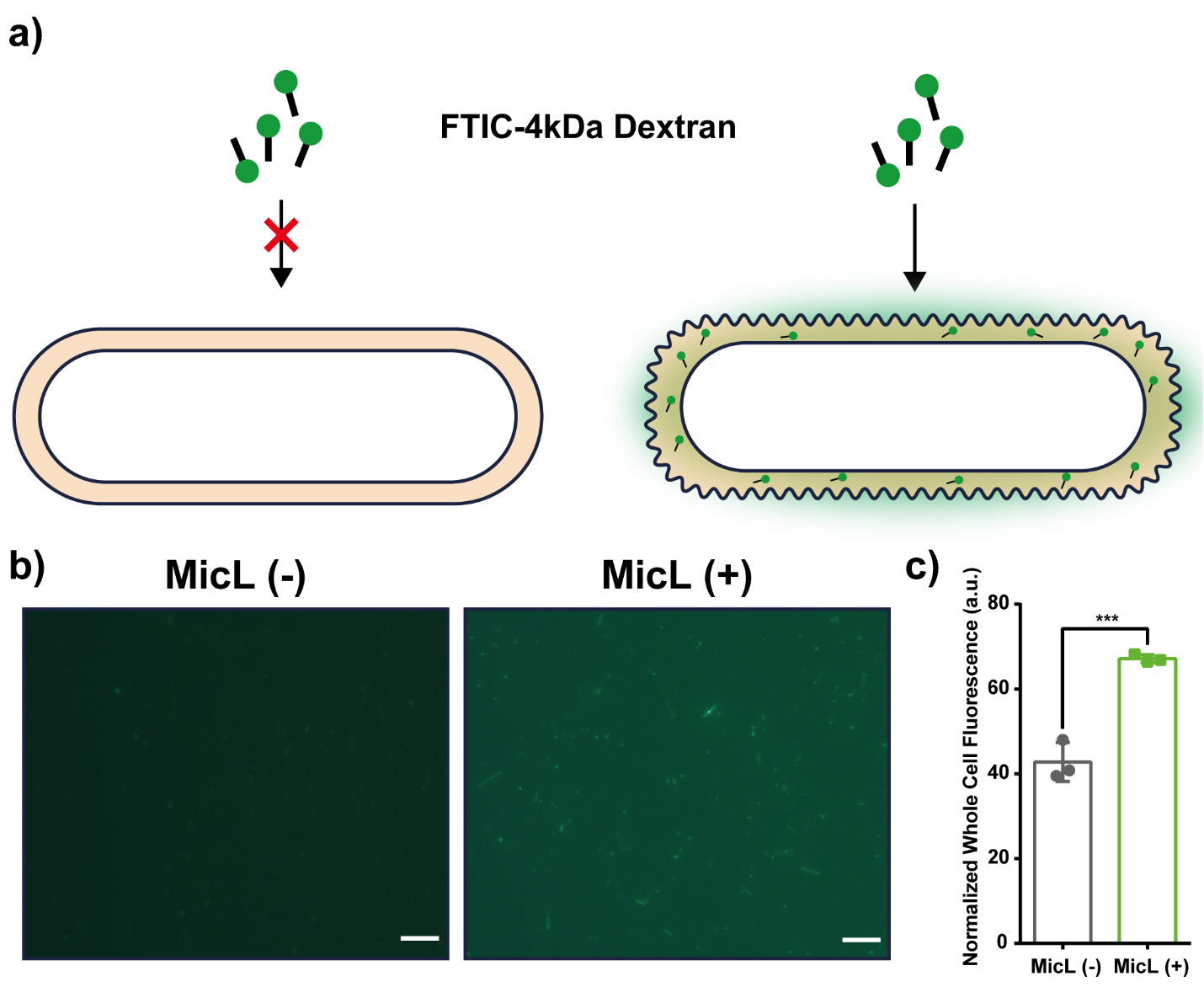

**Figure S1. iLOM enables diffusion of macromolecules form extracellular media to cell periplasm.** a) Schematic presentation of the assay to validate macromolecule diffusion from extracellular space to periplasm. b) Fluorescence microscope images of cells that treated with 4kDa Dextran conjugated with FTIC. Left; EcN without expression of micL sRNA, right; EcN cells expressing the micL sRNA c) Whole cell green fluorescence of cells with and without micL sRNA expression following to the diffusion assay.

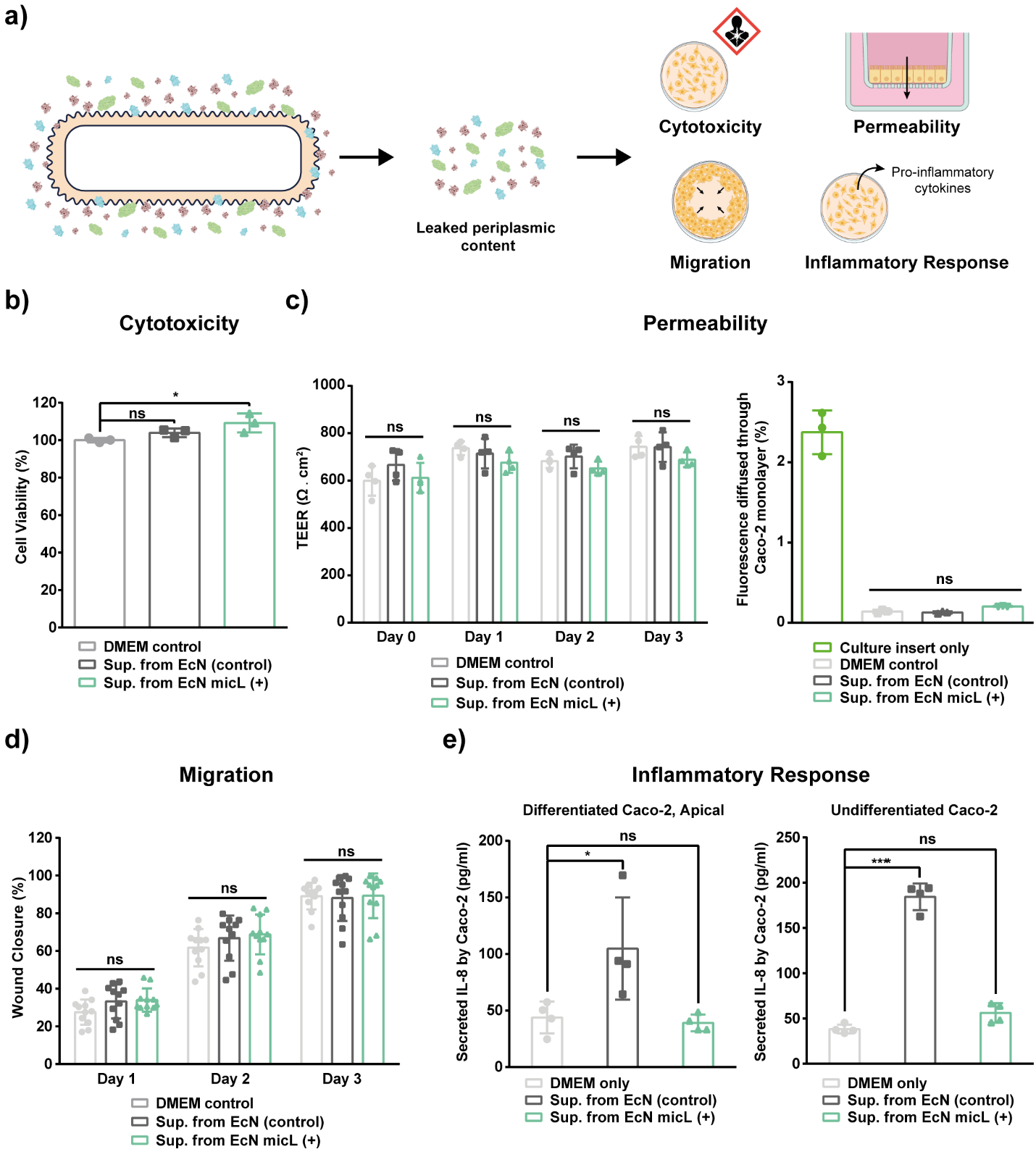

**Figure S2. *In vitro* safety assessments of iLOM-based secretion with EcN**. a) Schematic presentation of parameters that were tested with Caco-2 cell line. During the iLOM-based secretion, periplasmic content including proteins and other metabolites leaks through to extracellular space along with the PoI. b) Cytotoxicity of leaked periplasmic content from EcN. The experiment was conducted with three independent replicates c) Effects of leaked periplasmic content on permeability of differentiated Caco-2 monolayer. TEER (right) and FTIC conjugated dextran (kDa) diffusion (left) were determined to assess the permeability. TEER measurement was performed with four replicates while diffusion assay was done with three replicates. d) Wound closure assay was conducted to analyze the cell migration upon treatment with leaked periplasmic content. Percentage closure was calculated to normalize the data. Eleven replicates were used in the migration experiment. e) IL-8 secretion by Caco-2 cells treated with leaked periplasmic content. IL-8 amount was determined for undifferentiated (left) and apical side of differentiated (right) Caco-2 cells with ELISA. Four biological replicate was used.

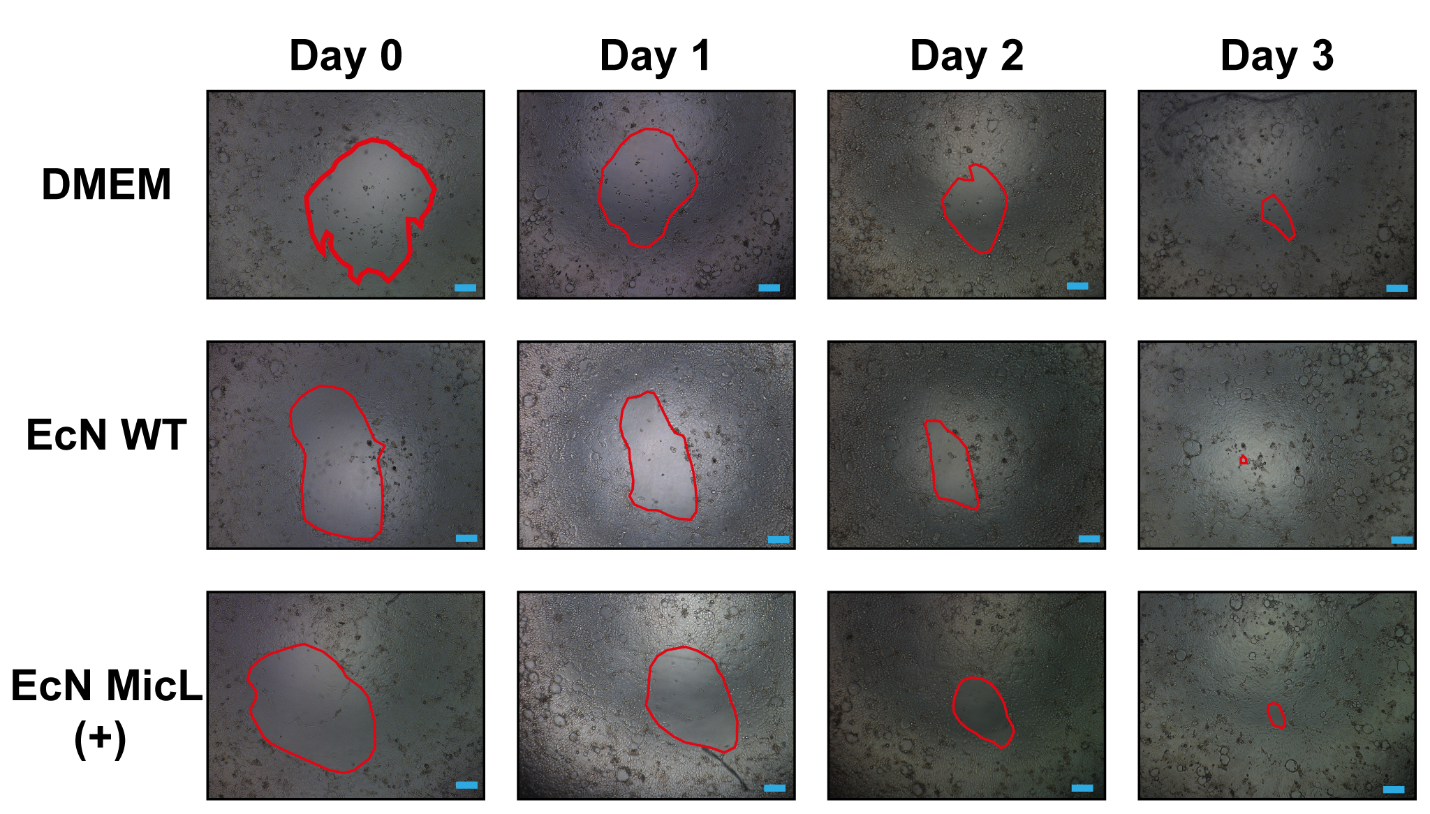

**Figure S3. Microscope images of holes in the migration assay.** Holes were imaged in each day for three days. For each image, the area of hole is calculated using ImageJ software.

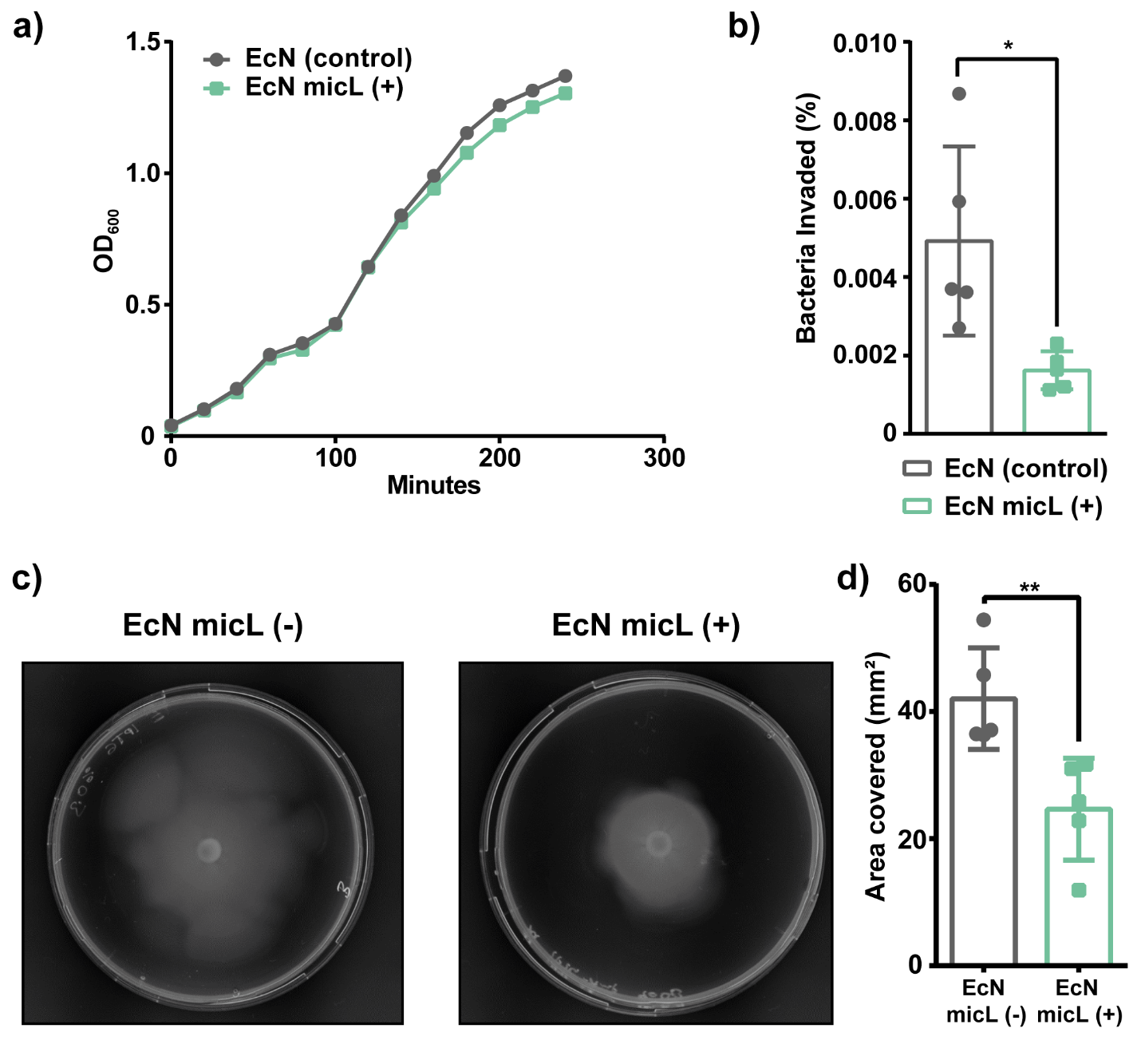

**Figure S4. Effects of micL sRNA on EcN physiology**. a) Growth curve of WT and micL sRNA expressing EcN. Optical density at 600nm was measured to monitor growth for 4 hours. Three biological replicates were used for the growth assay. b) Caco-2 invasion assay with WT and micL sRNA expressing EcN strains. The experiment was conducted with five biological replicates. c) Motility of WT and micL sRNA expressing EcN strains. The experiment was performed with five biological replicates.

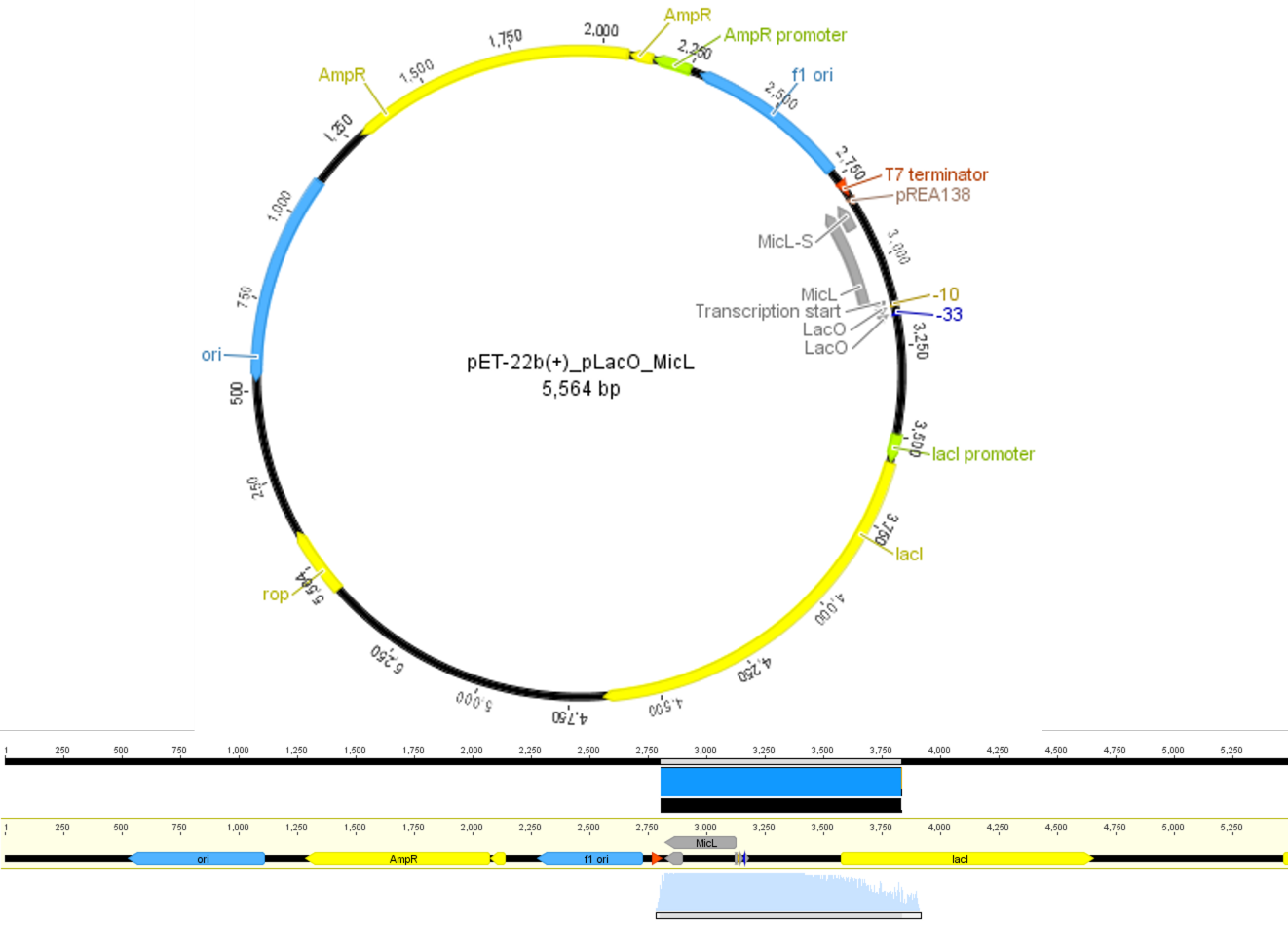

**Figure S5. Plasmid map of IPTG inducible micL sRNA in pET22b (+) vector and sequence verification of the vector with sanger sequencing.**

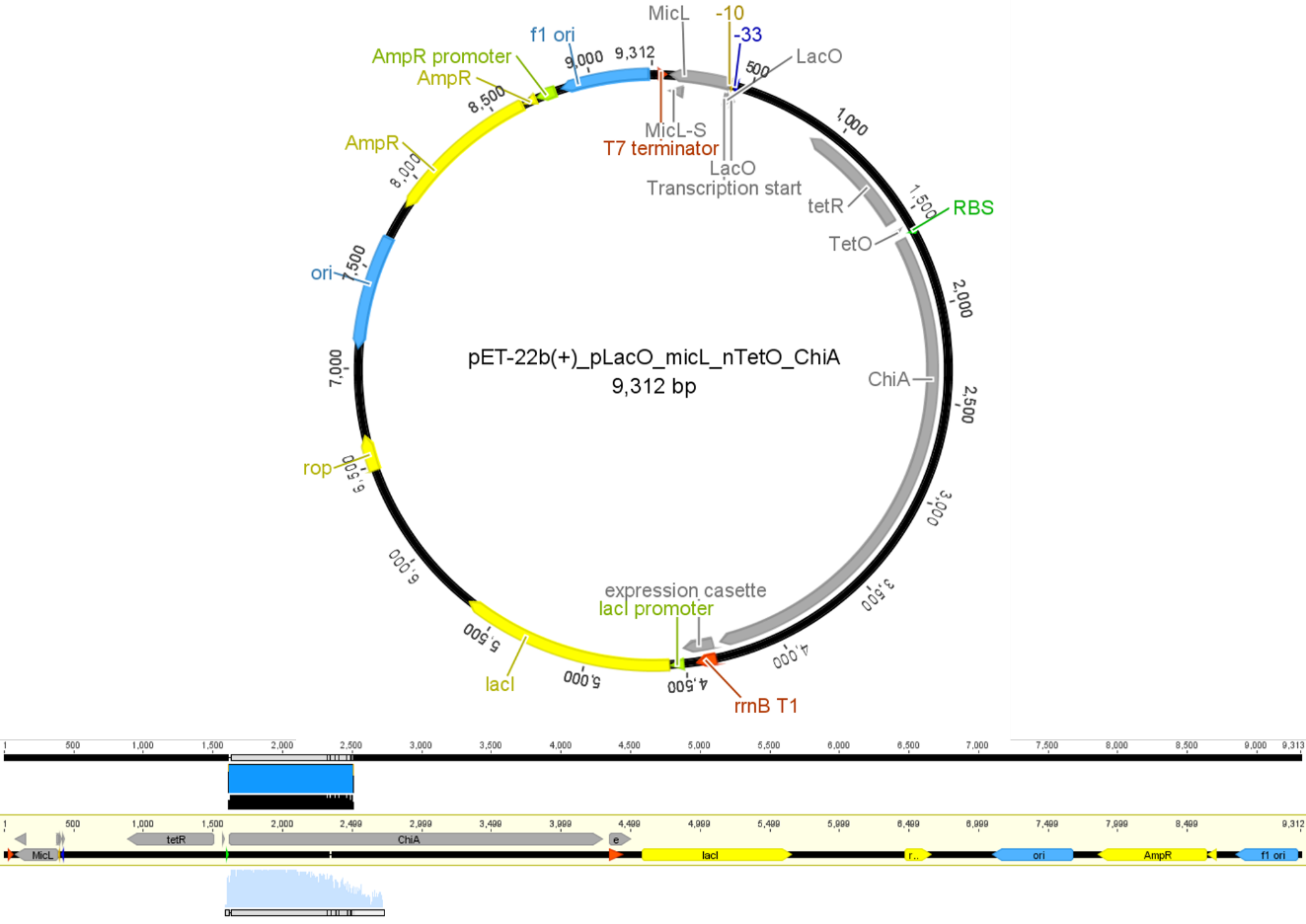

**Figure S6. Plasmid map of IPTG inducible micL sRNA and aTc inducible ChiA gene in pET22b (+) vector and sequence verification of the vector with sanger sequencing.**

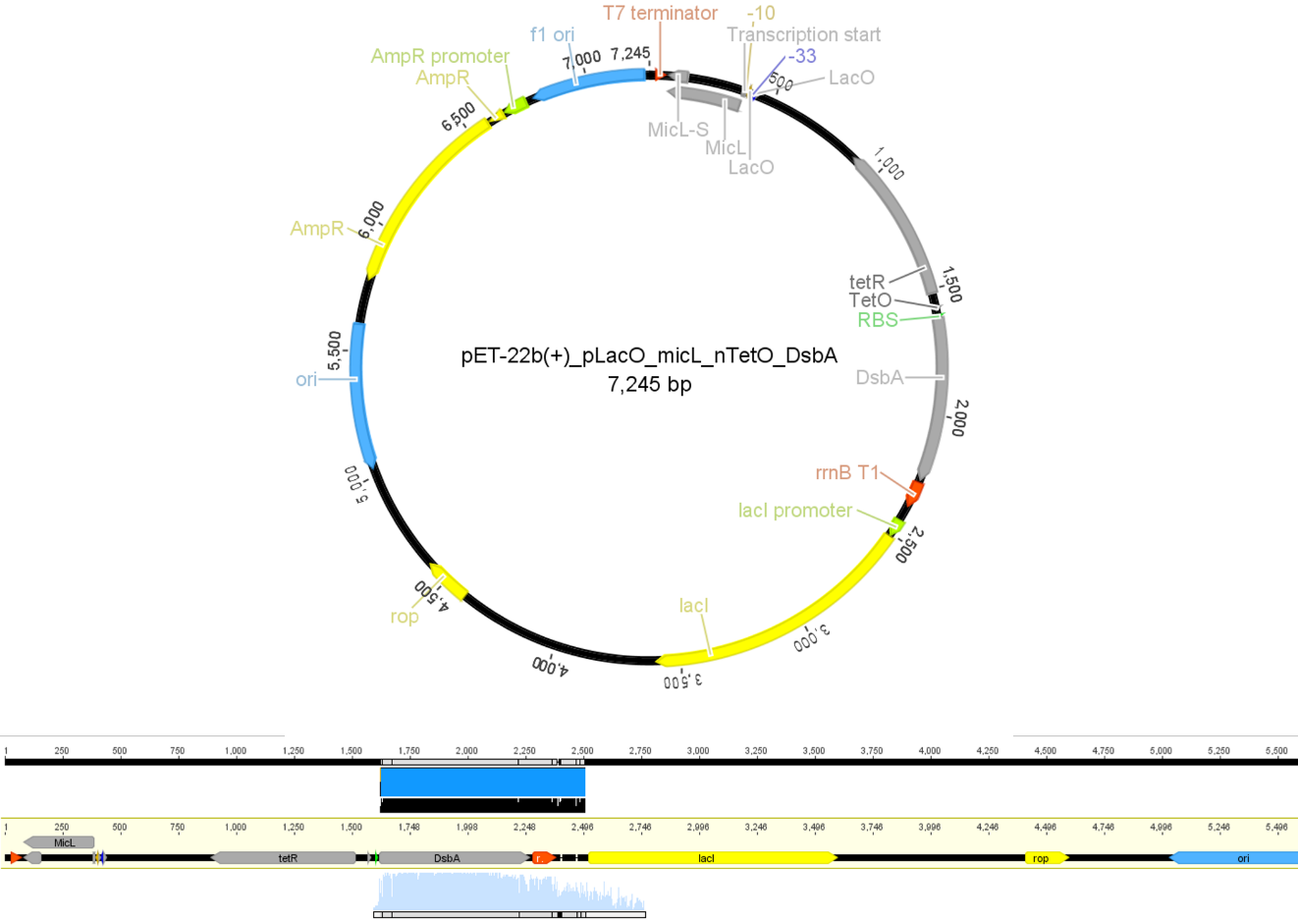

**Figure S7. Plasmid map of IPTG inducible micL sRNA and aTc inducible DsbA gene in pET22b (+) vector and sequence verification of the vector with sanger sequencing.**

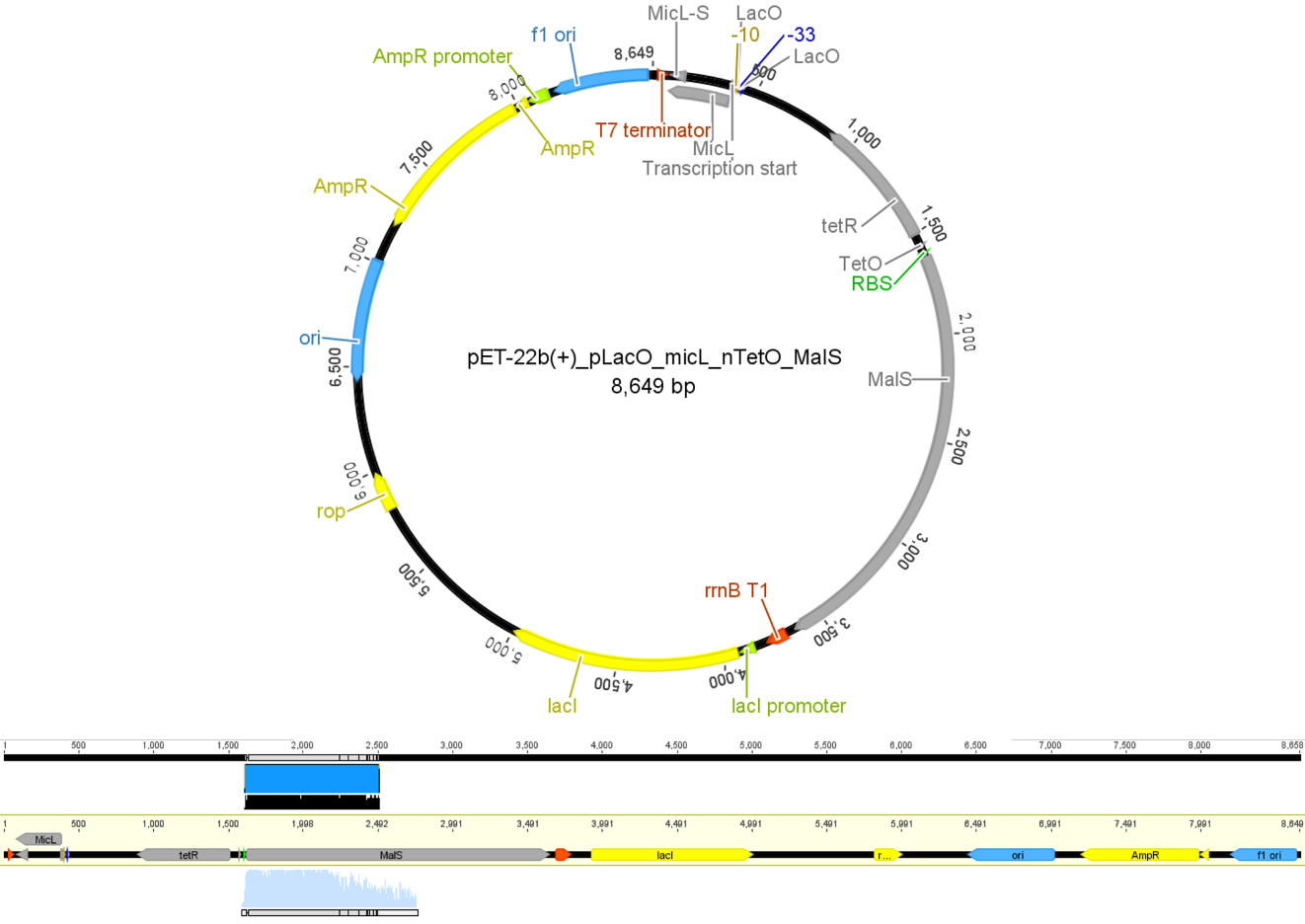

**Figure S8. Plasmid map of IPTG inducible micL sRNA and aTc inducible MalS gene in pET22b (+) vector and sequence verification of the vector with sanger sequencing.**

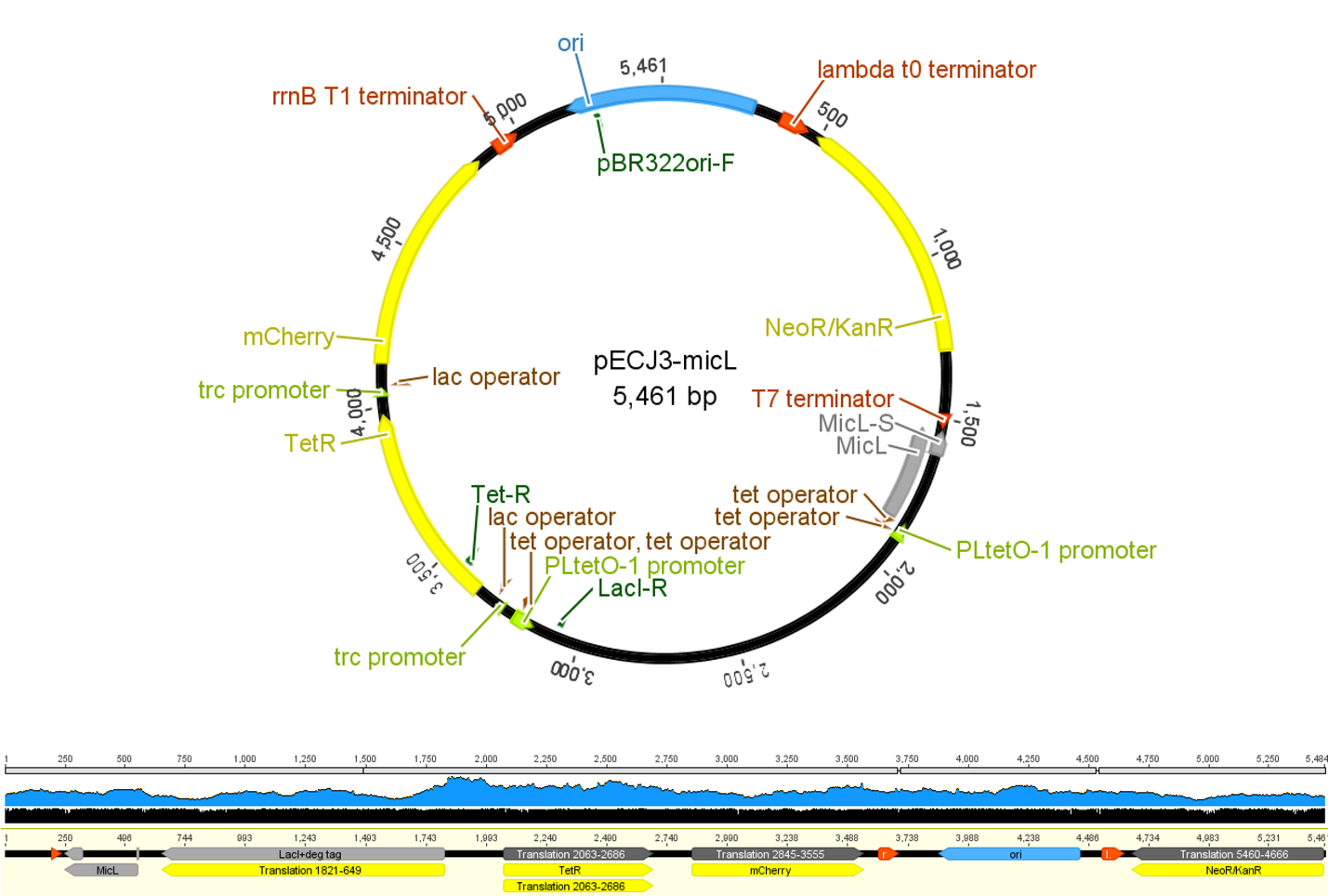

**Figure S9. Plasmid map of toggle switch circuit with micL in pECJ3 vector (ref) and sequence verification of the vector with NGS.**

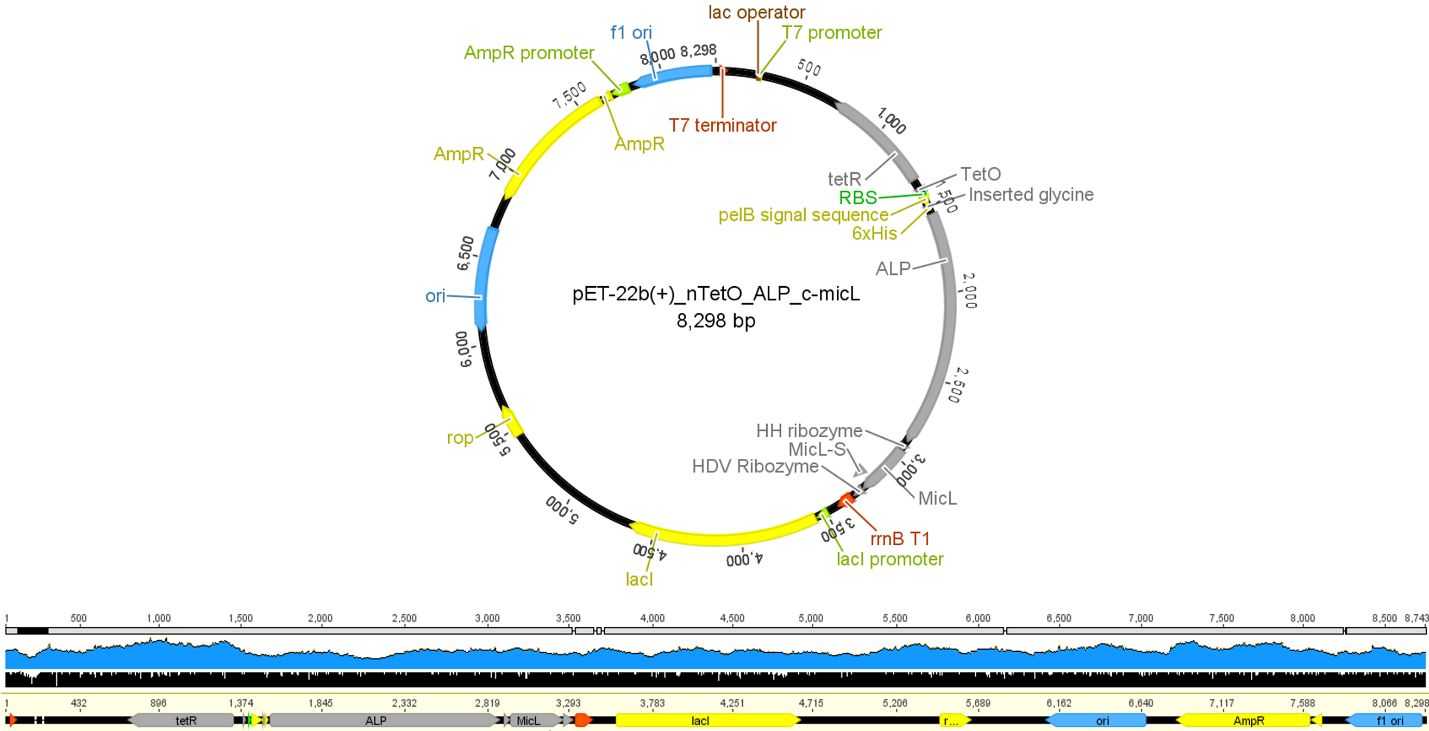

**Figure S10. Plasmid map of aTc inducible ALP (PhoA) gene fused with c-micL in pET22b (+) vector and sequence verification of the vector with NGS.**

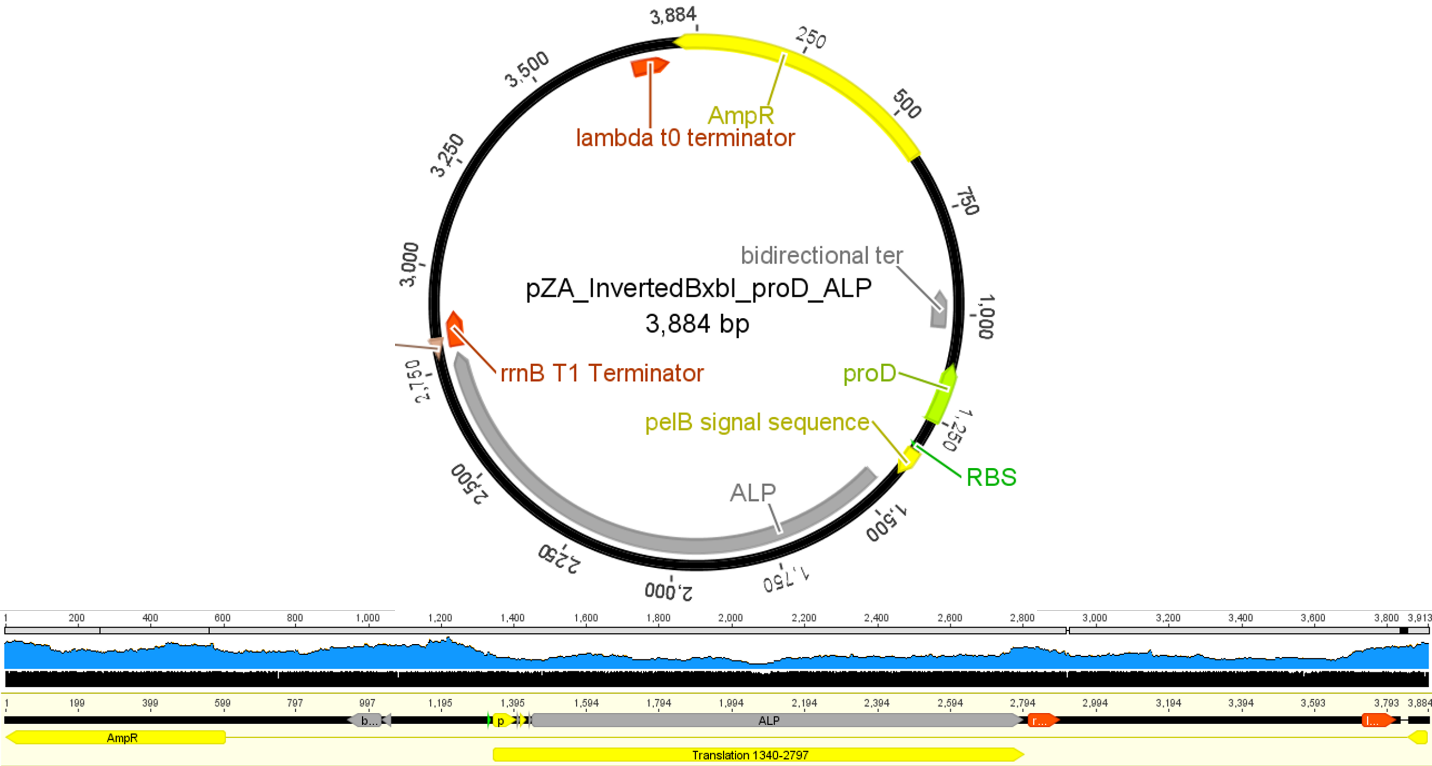

**Figure S11. Plasmid map of ALP (PhoA) gene located at inverted proD flanked by attB and attP sites of BxbI and sequence verification of the vector with NGS.**

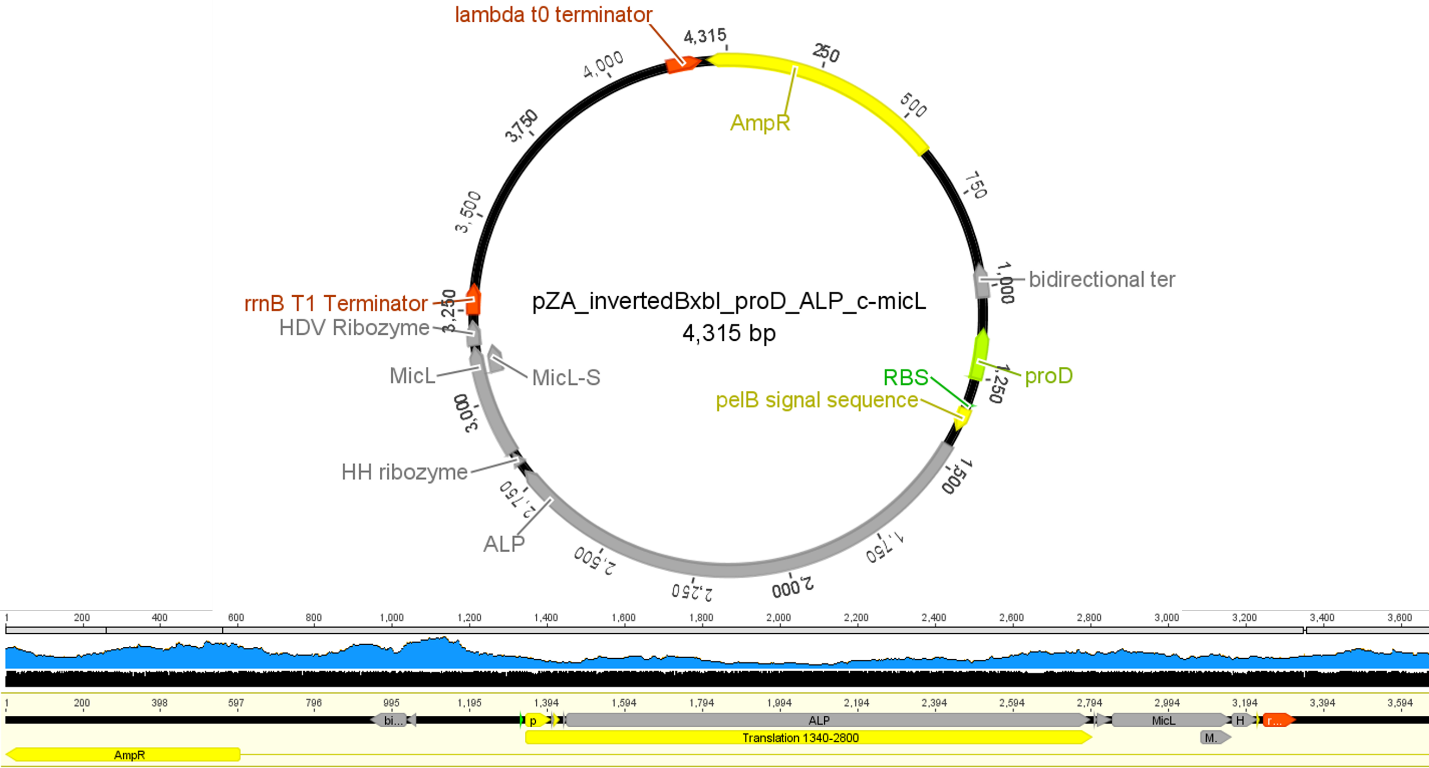

**Figure S12. Plasmid map of ALP (PhoA) gene fused with c-micL located at inverted proD flanked by attB and attP sites of BxbI and sequence verification of the vector with NGS.**

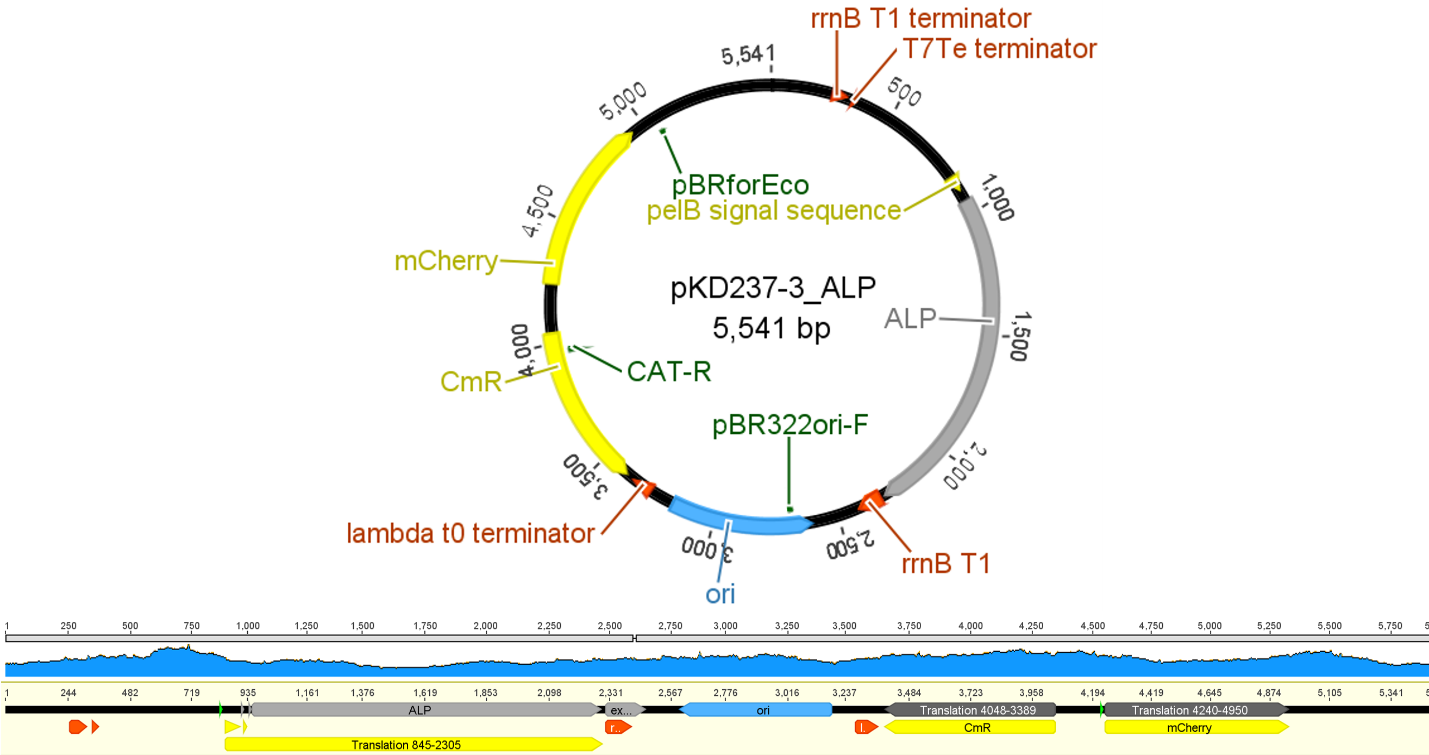

**Figure S13. Plasmid map of ALP (PhoA) cloned at the downstream of phsA in pKD237 vector (ref) and sequence verification of the vector with NGS.**

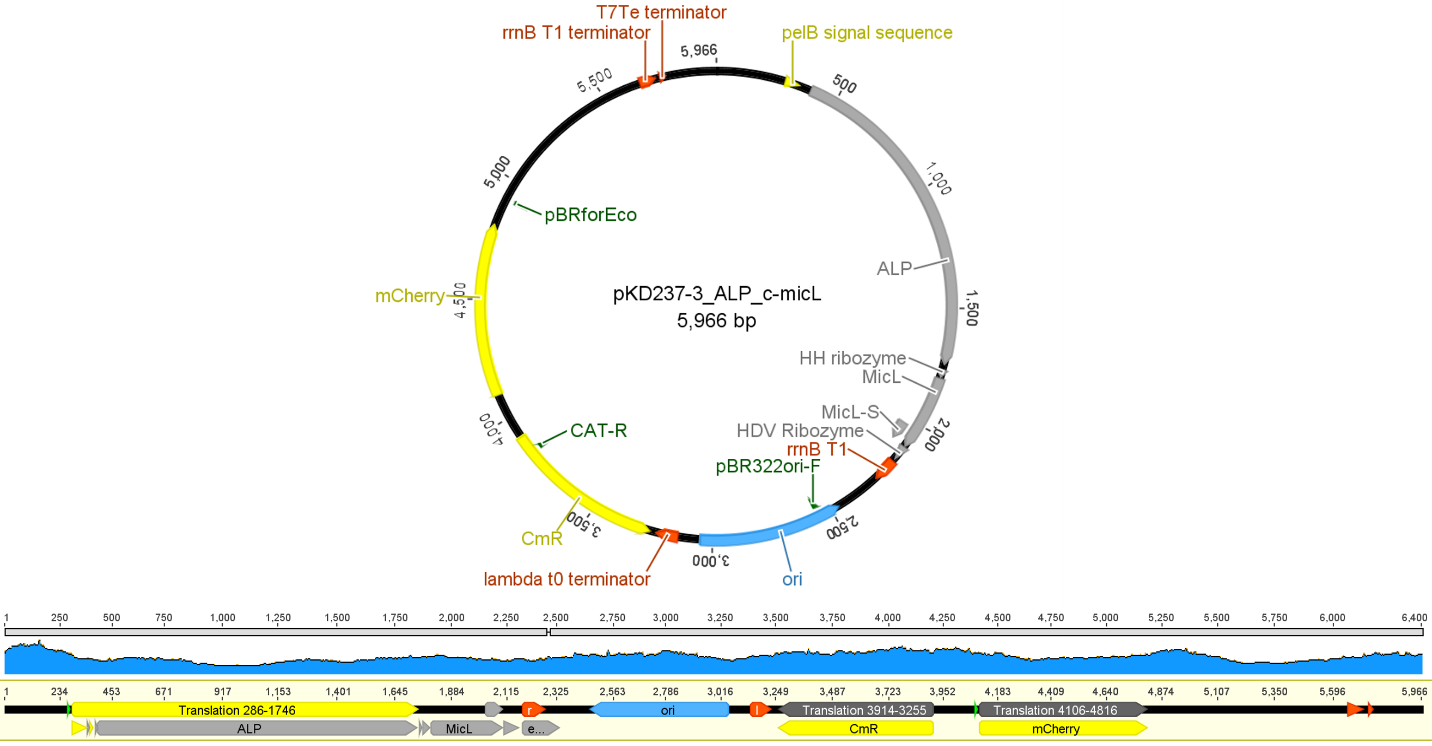

**Figure S14. Plasmid map of ALP (PhoA) fused with c-micL cloned at the downstream of phsA in pKD237 vector (ref) and sequence verification of the vector with NGS.**

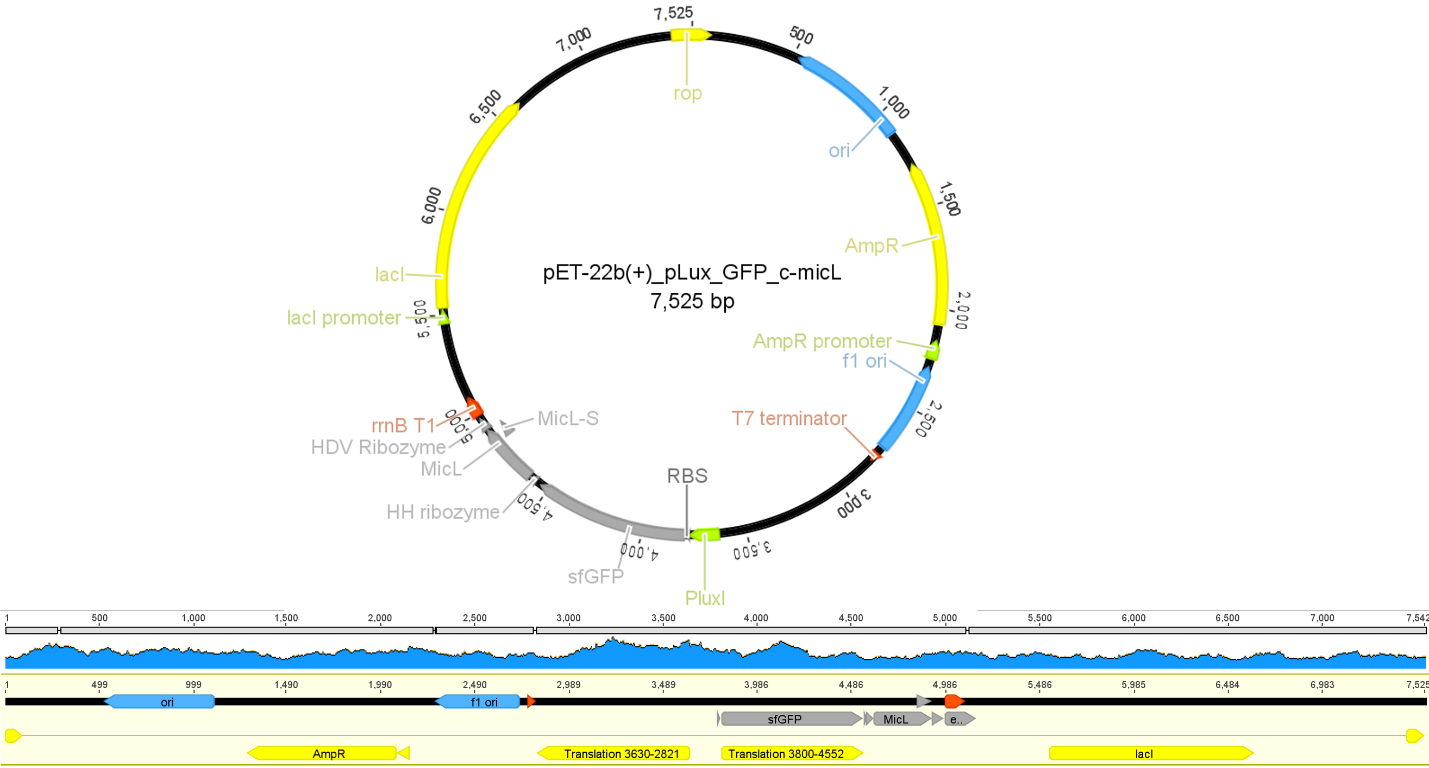

**Figure S15. Plasmid map of GFP fused with c-micL cloned downstream of pLux (ref) in pET22b (+) vector and sequence verification of the vector with NGS**

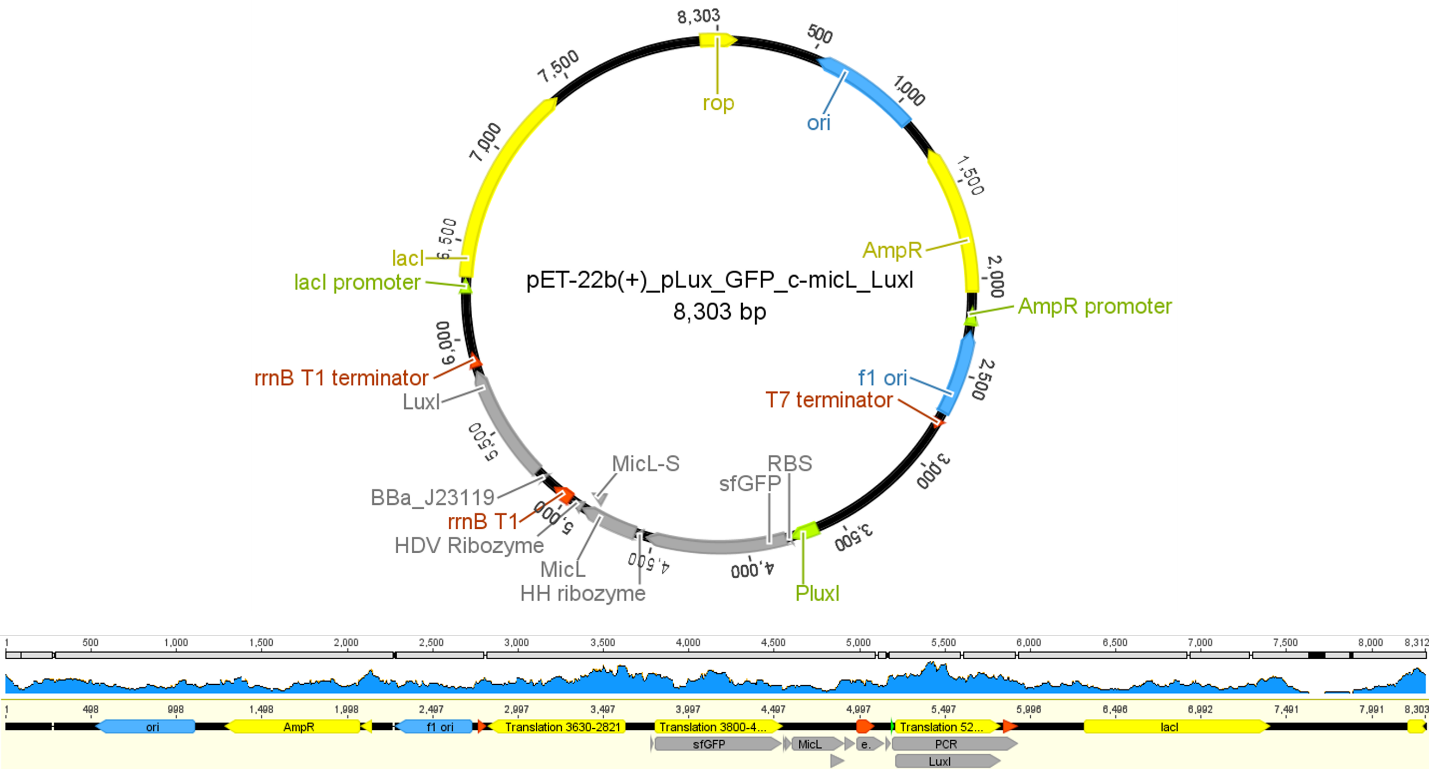

**Figure S16. Plasmid map of GFP fused with c-micL cloned at the downstream of pLux (ref) and constitutively expressed LuxI in pET22b (+) vector and sequence verification of the vector with NGS.**

.

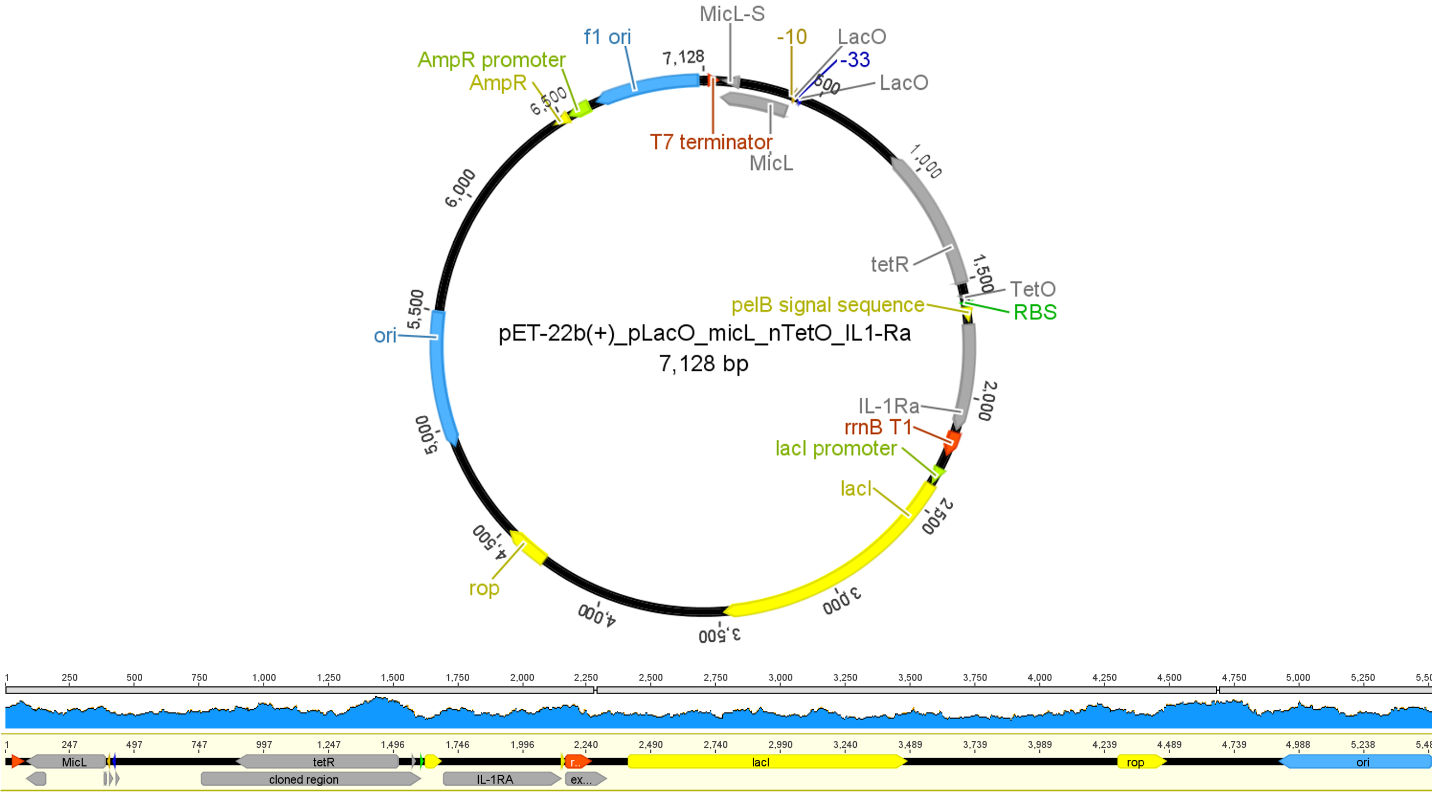

**Figure S17. Plasmid map of IPTG inducible micL sRNA and aTc inducible IL-1Ra gene in pET22b (+) vector and sequence verification of the vector with NGS.**

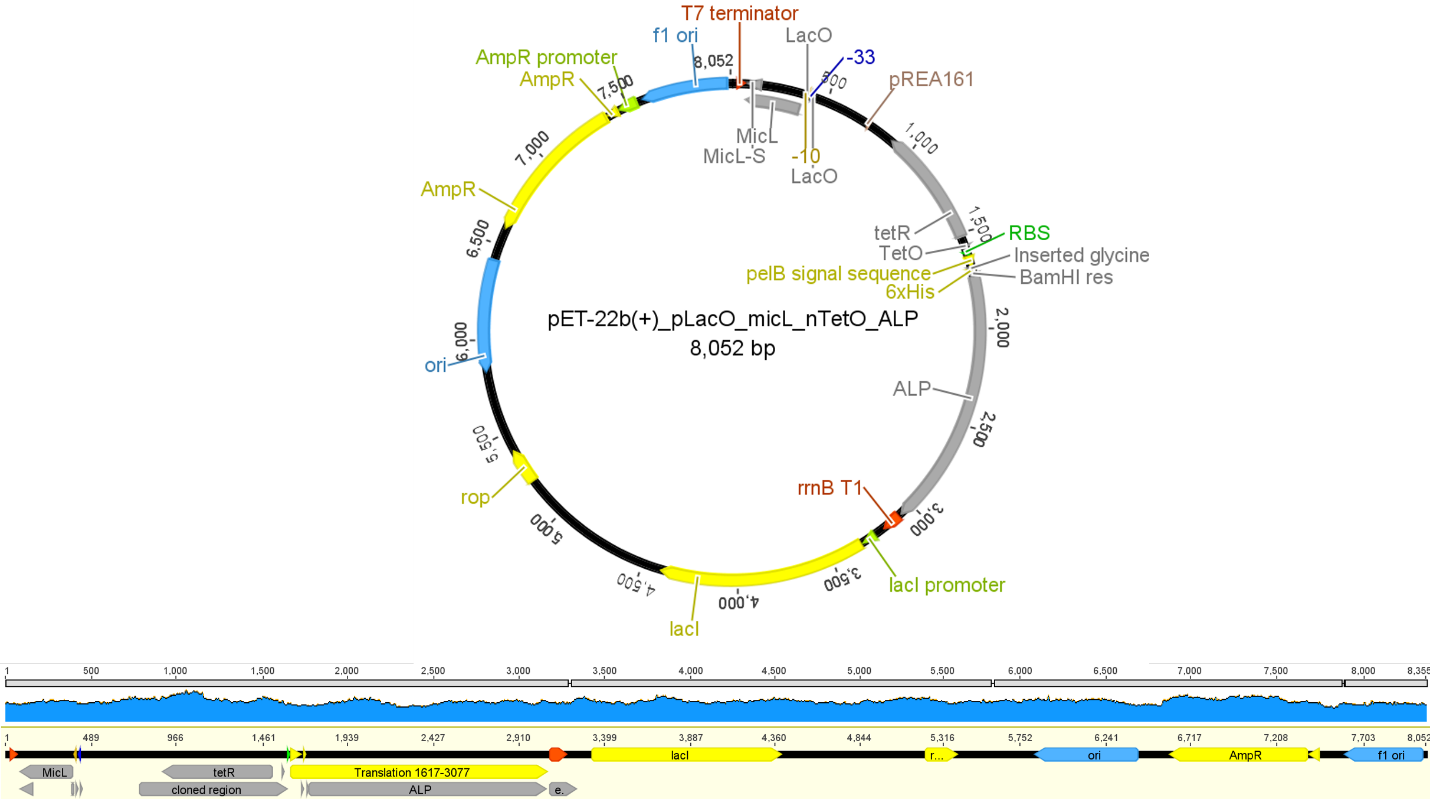

**Figure S18. Plasmid map of IPTG inducible micL sRNA and aTc inducible ALP (PhoA) gene in pET22b (+) vector and sequence verification of the vector with NGS.**

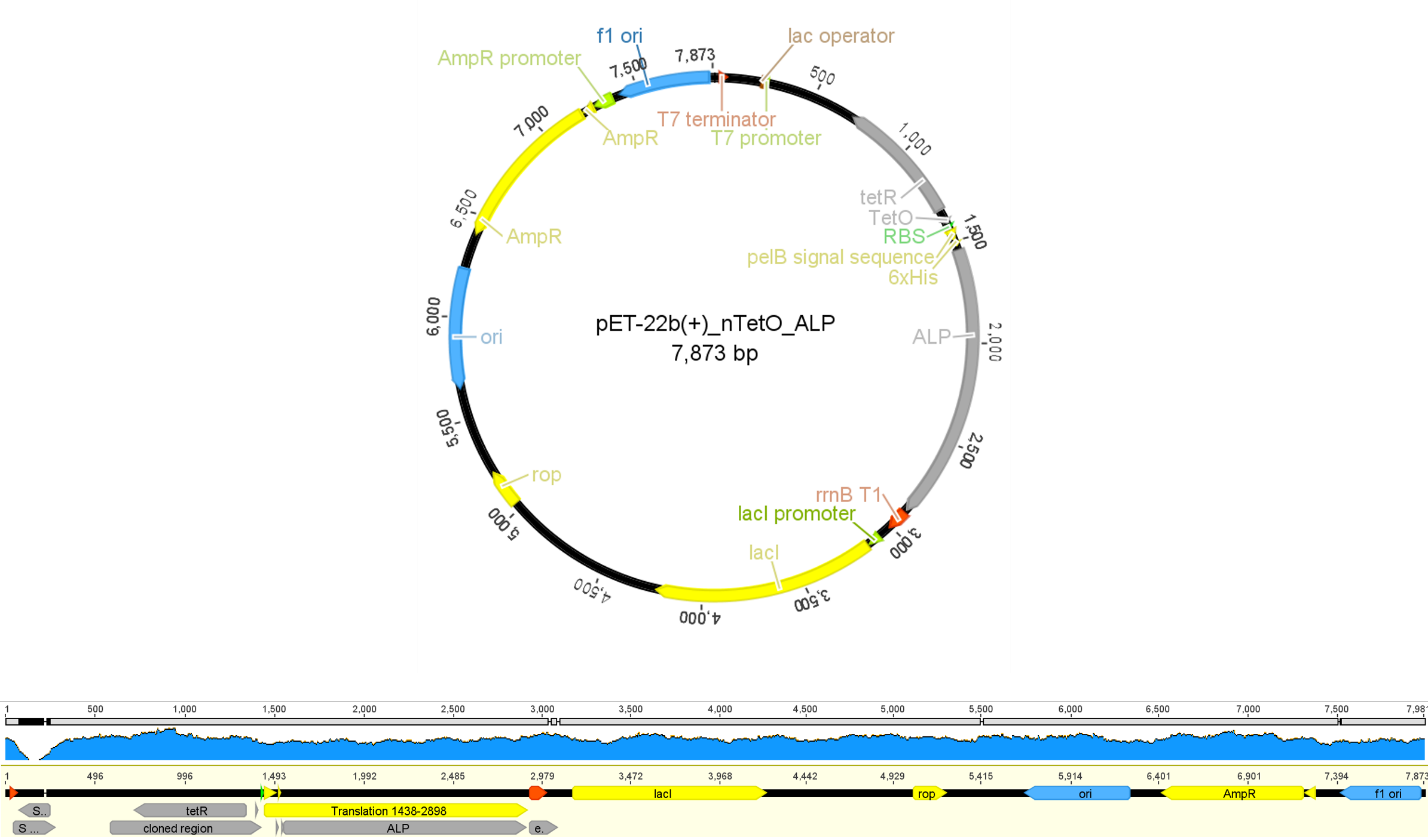

**Figure S19. Plasmid map of aTc inducible ALP (PhoA) gene in pET22b (+) vector and sequence verification of the vector with NGS.**

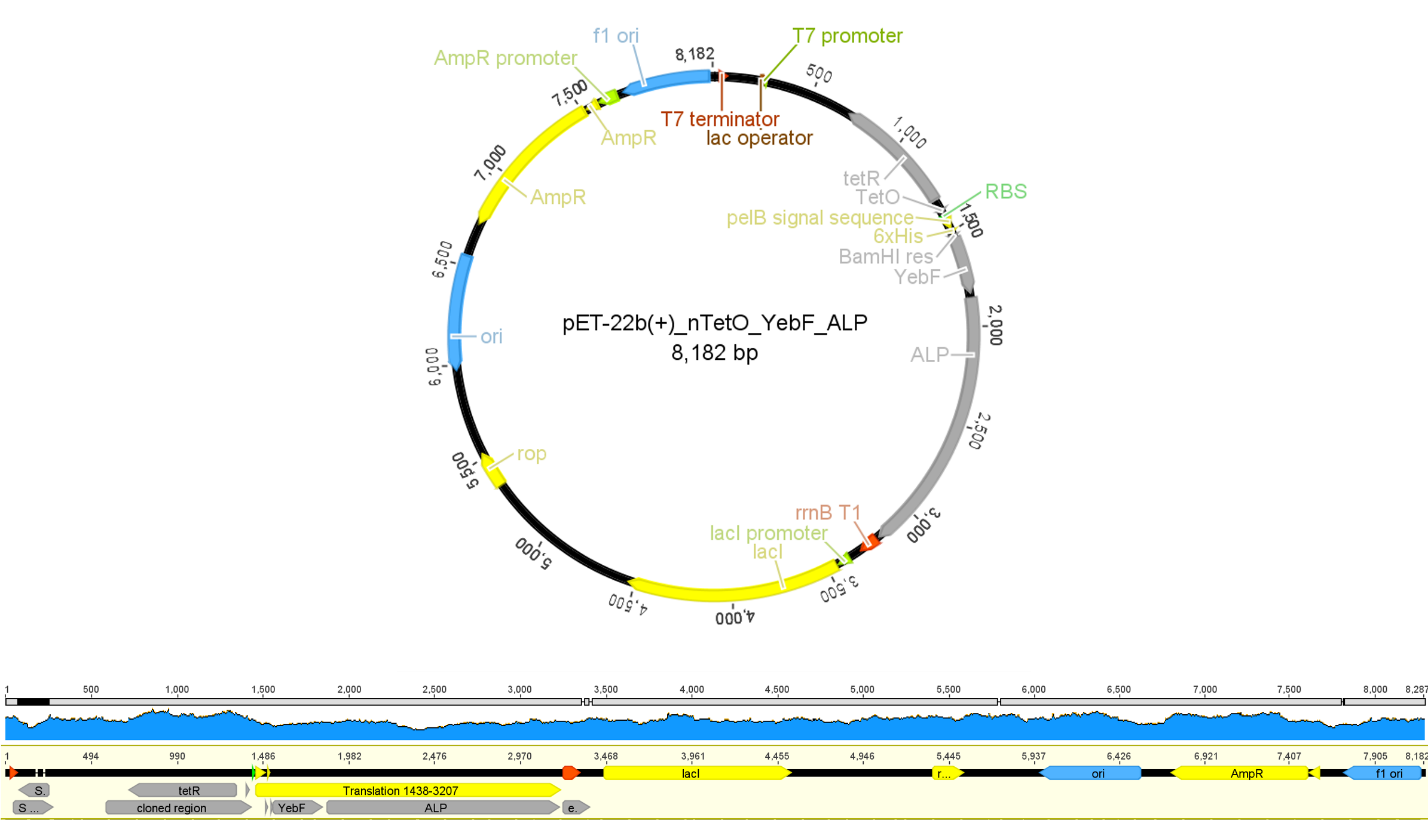

**Figure S20. Plasmid map of aTc inducible ALP (PhoA) gene fused with YebF gene in pET22b (+) vector and sequence verification of the vector with NGS.**
